# Residual avoidance: a new consistent and repeatable readout of chronic stress-induced conflict anxiety reversible by antidepressant treatment

**DOI:** 10.1101/414029

**Authors:** Thomas D. Prevot, Keith A. Misquitta, Corey Fee, Dwight F. Newton, Dipashree Chatterjee, Yuliya S. Nikolova, Etienne Sibille, Mounira Banasr

## Abstract

Stress-related illnesses such as major depressive and anxiety disorders are characterized by maladaptive responses to stressful life events. Chronic stress-based animal models have provided critical insight into the understanding of these responses. Currently available assays measuring chronic stress-induced behavioral states in mice are limited in their design (short, not repeatable, sensitive to experimenter-bias) and often inconsistent. Using the Noldus PhenoTyper apparatus, we identified a new readout that repeatedly assesses behavioral changes induced by chronic stress in two mouse models i.e. chronic restraint stress (CRS) and chronic unpredictable mild stress (UCMS). The PhenoTyper test consists of overnight monitoring of animals’ behavior in home-cage setting before, during and after a 1hr light challenge applied over a designated food zone. We tested the reproducibility and reliability of the PhenoTyper test in assessing the effects of chronic stress exposure, and compared outcomes with commonly-used tests. While chronic stress induced heterogeneous profiles in classical tests, CRS- and UCMS-exposed mice showed a very consistent response in the PhenoTyper test. Indeed, CRS and UCMS mice continue avoiding the lit zone in favor of the shelter zone. This “residual avoidance” after the light challenge, lasted for hours beyond termination of the challenge, was not observed after acute stress and was consistently found throughout stress exposure in both models. Chronic stress-induced residual avoidance was alleviated by chronic imipramine treatment but not acute diazepam administration. This behavioral index should be instrumental for studies aiming to better understand the trajectory of chronic stress-induced deficits and potentially screen novel anxiolytics and antidepressants.

## 1. Introduction

Depression is among the most debilitating disorders in the world (Friedrich, 2017) and commonly co-occurs with anxiety disorders (Hirschfeld, 2001). The proportion of the global population living with depression and anxiety recently reached 4.4% and 3.6%, respectively (WHO, 2017) with ~85-90% of diagnosed patients suffering from both conditions (Tiller, 2013). Human studies have been useful to identify biological deficits associated with these disorders, and since the early 1980’s, great progress has been made in elucidating pathological mechanisms and exploring new avenues for therapeutic interventions through using animal models (Cryan and Sweeney, 2011; Katz and Hersh, 1981; Katz et al., 1981).

The most popular rodent models employ exposure to chronic stress to study the psychopathology of mood disorders including depression (Chiba et al., 2012; Eiland and McEwen, 2012; Vyas et al., 2004; Willner, 2017a). The unpredictable chronic mild stress (**UCMS**) model has strong face and construct validities for modeling in rodents behavioral dimensions relevant to human stress-related illnesses (Willner, 1997, 2017a). This model as well as the chronic restraint stress (**CRS**) or immobilisation stress paradigms are often used to study cellular mechanisms of stress-related disorders, including changes in neuronal spine density, cell signaling, or neurotransmitter systems which parallel human cellular pathologies associated with major depressive disorders (Kang et al., 2012; McEwen, 1997; McEwen and Magarinos, 1997; Qiao et al., 2016). At the behavioural level, past studies demonstrate that exposing mice or rats to chronic stress paradigms leads to persistent cognitive deficits (Sandi, 2004), helplessness- (Strekalova and Steinbusch, 2010), anhedonia- (Willner, 2017a), and anxiety-like (Belzung and Lemoine, 2011) behaviors. However, these findings have been criticized for being difficult to replicate (Castanheira et al., 2018; Cryan and Sweeney, 2011; Ferreira et al., 2018; Willner, 2017a, b). Recently, a survey of experimenters using UCMS paradigms found that 21% experienced difficulties replicating expected results (Willner, 2017b). The lack of reliability, or the inability to induce expected phenotypes, is often attributed to individual variability (e.g., strain, sex, or rearing conditions) or non-optimal methods for assessing chronic stress-induced phenotypes. Nevertheless, some responsibility for this lack of reproducibility can be attributed to the behavioral tests used. A vast majority of behavioral tests employed to study the effects of chronic stress measure anxiety-like behaviors, and were primarily designed to detect anxiolytic and antidepressant drug efficacy in non-stress animals (Bruhwyler et al., 1995; Cryan et al., 2002; Surget et al., 2008). These tests are also limited by protocol variability within and across labs, experimenter bias, and the requirement for novelty precluding repetitive measurement over chronic time courses. More precisely, there are currently >15 different assays regularly employed to assess anxiety-like behavior and adapted to investigate the effects of chronic stress in rodents (Ohl, 2005). Most are relatively short (5-30 minutes) and based on approach-avoidance (Bailey and Crawley, 2009), i.e. conflict between animals’ innate exploratory drive and their aversion to a threatening environment (Rodgers and Dalvi, 1997). In addition, most of these behavioral tests have a novelty component which limits their ability to provide a time-dependent trajectory of phenotype development, maintenance, or reversal that can be of primary importance in studying chronic stress.

For a better understanding of the dynamic progression of behavioral deficits induced by chronic stress, it is necessary to develop experimenter-free, repeatable, reliable, and effective tests measuring chronic stress-induced behavior in rodents. Here, we first aimed to demonstrate the limitations of commonly-used tests using two paradigms in mice, UCMS and CRS. Then, we set out to demonstrate the usefulness of the PhenoTyper test, a recently designed test that can assess chronic stress-induced conflict anxiety repetitively with minimal experimenter bias, and is based on measuring the animal’s activity in response to an aversive light challenge in a home-cage like setting (Nikolova et al., 2018; Maluach et al., 2017; Aarts et al., 2015). In depth analysis of the effects of chronic stress exposure (weekly characterization and cross models comparison) revealed a highly reproducible chronic stress-specific pattern of response after the light challenge, where the animals continue to avoid the lit zone. Here, we develop a new calculation/readout which we named “residual avoidance” to establish a quantifiable measure of this behavior, and then demonstrated its ability to evaluate the behavioral trajectories of chronic stress and antidepressant treatment.

## 2. Material and Methods

### 2.1. Animals

Eight-week old C57B6 mice (Jackson Laboratory, Bar Harbor, Maine, USA; 50%♀) were housed under normal conditions with a 12hr light/dark cycle and provided with *ad libitum* access to food and water. All animals were kept for 1-2 weeks in the facility before the start of the experiments. Experimenters were blind to treatment group assignments during behavioral testing. All procedures followed guidelines set by the Canadian Council on Animal Care (CCAC). Six cohorts were used in this study. One cohort was used to assess baseline behavior and light challenge response in the PhenoTyper apparatus (n=12/group, 50% females). The second experiment measured the effects of acute restraint stress (**ARS**) in this test (n=8/group, 50% females). In Experiment 3, we compared the effects of group-housing (3 mice per cage) vs. single-housing for 5 weeks and only tested the animals in PhenoTyper once at the end of the experiment (n=12/group, 50% females). Experiments 4 and 5 tested the effects of CRS and UCMS respectively and, the last two cohorts were used to assess the efficacy of acute diazepam (anxiolytic) or chronic imipramine (antidepressant) treatment in CRS animals.

#### 2.1.1. Unpredictable chronic stress (UCMS)

UCMS mice were subjected to randomized stressors (3-4/day) for 5 weeks (Nikolova et al., 2018). These stressors included: forced bath (~2cm of water in the cage for 15min), wet bedding, aversive smell (20min to 1hr exposure to fox or bobcat urine), light cycle reversal or disruption, bedding exchange (rotate mice into previously occupied cages), tilted cage (45° angle), reduced space, restraint (50mL falcon tube with nose/tail holes for 15-30min), bedding change (replacing soiled with clean bedding), no bedding, or nestlet removal. UCMS animals are single housed throughout the experiment as part of the UCMS procedure. Control animals were group-housed (~3-4/cage) to eliminate the stress of single-housing (n=12/group, 50% females).

#### 2.1.2. Restraint stress

Animals were placed into 50mL falcon tubes with nose and tail holes for air flow. ARS consists in one session of 1hr before testing. For CRS, restraint sessions occurred twice daily for 1hr during the light cycle (7:00am-7:00pm), separated by a minimum of 2hrs. CRS-exposed animals were single-housed throughout the stress exposure and control animals were group-housed (n=12/group, 50% females). One CRS male mouse died during the experiment and was removed from the study.

#### 2.1.3. Drug administration

After 3 weeks of CRS exposure, mice (n=10-12/group, 50% females) received diazepam (1.5 mg/kg i.p.) or vehicle (0.9% saline) acutely 30min before being placed in the PhenoTyper or NSF arenas. Tests were performed 48hrs apart. To assess if CRS effects were reversible by chronic antidepressant treatment, mice subjected to CRS for 3 weeks received imipramine (~15mg/kg in drinking water based on ~8mL/day fluid consumption) for the 3 following weeks while continuing CRS exposure (n=12/group, 50% females). Freshly made imipramine was provided every other day to prevent deterioration due to light exposure or room temperature. Animals were tested weekly in the PhenoTyper test and once in the NSF, 48hrs after the last PhenoTyper test.

### 2.2. Body weight and coat state

Body weight and fur coat deterioration of mice were assessed weekly. Weight gain was calculated using baseline weights (week 0, before chronic stress exposure), and expressed in percentage of change. Fur coat deterioration was measured using a rating scale of 0-1, with 0 equivalent to a well-groomed, smooth coat and 1 equivalent to a tousled or soiled coat with bald patches (Nollet et al., 2013). Coat quality was assessed for 7 anatomical areas (head, forepaws, hindpaws, back, neck, tail, and abdomen) and added for a single score of coat state deterioration per animal.

### 2.3. Behavioral Tests

Following 5 weeks of stress exposure, mouse behavior was assessed in a series of commonly-used tests measuring chronic stress-induced behavioral phenotype: the elevated plus maze (EPM), open field (OF), novelty-induced hypophagia (NIH), and novelty-suppressed feeding (NSF) tests. Tests were spaced 48hrs apart to minimize between-test interaction. On non-testing days, CRS or UCMS were resumed to maintain stress-induced profiles. Tests were performed a minimum of 12hrs after the last stressor.

#### 2.3.1. EPM test

The EPM apparatus consist of four Plexiglas arms, where two open arms (67×7 cm) and two enclosed arms (67×7×17 cm) form a plus shape with similar arms facing each other. The EPM is situated 95cm above the floor within a dimly lit room (20 lux). Mice are placed in the center of the intersecting arms, facing an open arm, and are allowed to explore the apparatus for 10min. A digital camera mounted on a rod recorded animal behavior from above. *A posteriori* measurement of time spent and the number of entries into open and closed arms is performed using ANYmaze (Stoelting Co.™, Wood Dale, IL, USA).

#### 2.3.2. OF test

The OF apparatus consists of a 50×50 cm chamber that each mouse explores freely for 10min. Time spent and total entries into the center of the arena (digitally defined 20×20 cm zone in the center of the apparatus) were measured. A digital camera mounted on a rod recorded animal behavior from above. *A posteriori* measurements were performed using ANYmaze.

#### 2.3.3. NSF test

Following ~16hrs food deprivation, mice are placed in a novel arena (62×31 cm dimly lit enclosure) containing a single food pellet. Latency to feed on the pellet is measured for a 12min period. To control for appetite drive and/or food deprivation-induced weight loss, latency to feed was measured in the animal’s home cage following the novel environment test for a maximum of 6min.

#### 2.3.4. NIH test

Mice are habituated to a palatable liquid (1mL of 1:3 sweetened condensed milk in water) in a Petri dish for 3 consecutive days. On the following 2 days, mice are timed for their milk approach and drinking latency across 2 test sessions. The first session occurs in the home cage under normal light conditions and serves as a control for potential treatment effects on appetite or activity. Home cage latency to consume the milk from the Petri dish is recorded. 24hrs later, the subsequent test session occurs in a similar but new cage and under bright lighting (500-600 lux). Latency to consume measured during the second test (novel environment) is used as an index of hyponeophagia. Tests sessions occur over a maximum duration of 10mins.

#### 2.3.5. PhenoTyper™ test

The Noldus™ PhenoTyper apparatus (Leesburg, VA, USA) is a commercially-available observation cage designed for video-tracking of regular mouse behavior over extended periods using the Noldus EthoVision 10 software. The apparatus contains an integrated infrared sensitive camera that tracks activity and time spent by the animal in customizable zones. We established two designated zones (**Fig. S1**), a food zone (6.5×15 cm) and a shelter zone (10×10 cm), and tracked the animals’ location throughout the dark cycle (19:00-07:00). The apparatus also contains a white LED spotlight placed above the food zone, a feature that we employed to test whether animals would alter normal behavior in response to the application of an aversive “light challenge”. Specifically, we set the spotlight to turn on 4hrs into the dark cycle for a 1hr period (11:00pm-12:00am). This time-window was strategically chosen for being within a naturally high plateau for time spent in the food zone by the animal. For chronic stress studies, the PhenoTyper test was performed weekly on days 0, 7, 14, 21, 28, and 36. For all experiments, food and shelter zone time was measured in 1hr bins.

Based on the time spent in food and shelter zones after the light challenge, an index of avoidance was calculated as a function of the control group’s response to the light challenge. This calculation was named Residual Avoidance (RA) and considers the difference between the animals’ response during the light challenge and the sum of time spent avoiding the lit zone for the following 5hrs. The 5hrs cut-off was chosen to cover the time window of potential lasting response to the light challenge but exclude the last 3hrs of the dark cycle when animal’s time spent in shelter or food zones plateau to the maximum and minimum, respectively. RA was calculated for each mouse as followed:

**In the food zone:**

[1-(Σ Time_(12am-5am)_ – Time_(11pm-12am)_)/ Average control group (Σ Time _(12am-5am)_ – Time_(11pm-12am)_)] * 100, where “Time” is time spent if food zone.

**In the shelter zone:**

[(Σ Time_(12am-5am)_ – Time_(11pm-12am)_)/ Average control group (Σ Time _(12am-5am)_ – Time_(11pm-12am)_)-1] * 100, where “Time” is time spent if shelter zone.

Residual avoidance provides information about how animals react after the light challenge, relative to the designated control group. Control group animals have a mean RA=0, while positive RA means that an animal displays avoidance for either food or shelter zones from the zone light challenge.

### 2.4. Statistics

Statistical analyses were performed using Statview software (SAS Institute Inc., Cary, NC, USA). Student’s t-test was used to determine statistical differences between groups for commonly-used behavioral tests. Repeated-measures ANCOVA (analysis of covariance) was used for within- and between-subject evaluation of behavioral changes in the PhenoTyper test, and for time course analysis of longitudinal assessments including weight gain, coat state, or weekly RA. Sex was included as a covariate for all analyses. Stress and drug effects were assessed using repeated measures ANCOVA for RA data and two-way ANCOVA for NSF data. Significant results were followed-up with *post-hoc* Fisher’s test. In addition, summary scores capturing emotional reactivity dimensionally across groups were generated using principal component analysis (PCA) on 16 major behavioral variables collected throughout the last week of testing. Variables included all key behavioral parameters from commonly-used tests, food and shelter times before and during challenge as well as RAs for both zones. For longitudinal measures, the PCA included the last readout only. All sets of variables were submitted to PCA with a varimax rotation based on a correlation matrix with mice as one experimental unit. Two-way ANCOVA and *post-hoc* analysis was used to determine stress and sex contributions to the first 3 components. For results interpretation, the loadings of the individual variables onto each of the newly derived components were examined. PC loadings are expressed in absolute value, loadings >0.4 were considered significant. This cut-off was based on the critical r values corresponding to p<0.05 for the current sample size. PCA and follow up analysis was performed using SPSS software (IBSM SPSS statistic 24).

## 3. Results

### 3.1. Commonly-used behavior tests assessing chronic stress-induced behaviors revealed heterogeneous profiles between tests and models

Mice were exposed to CRS (**Figure 1 A-F**) or UCMS (**Figure 1 G-L**) for 5 weeks before behavioral assessment using the EPM, OF, NSF, and NIH tests. Coat state and weight gain were assessed before and throughout stress exposure.

**Figure 1:**
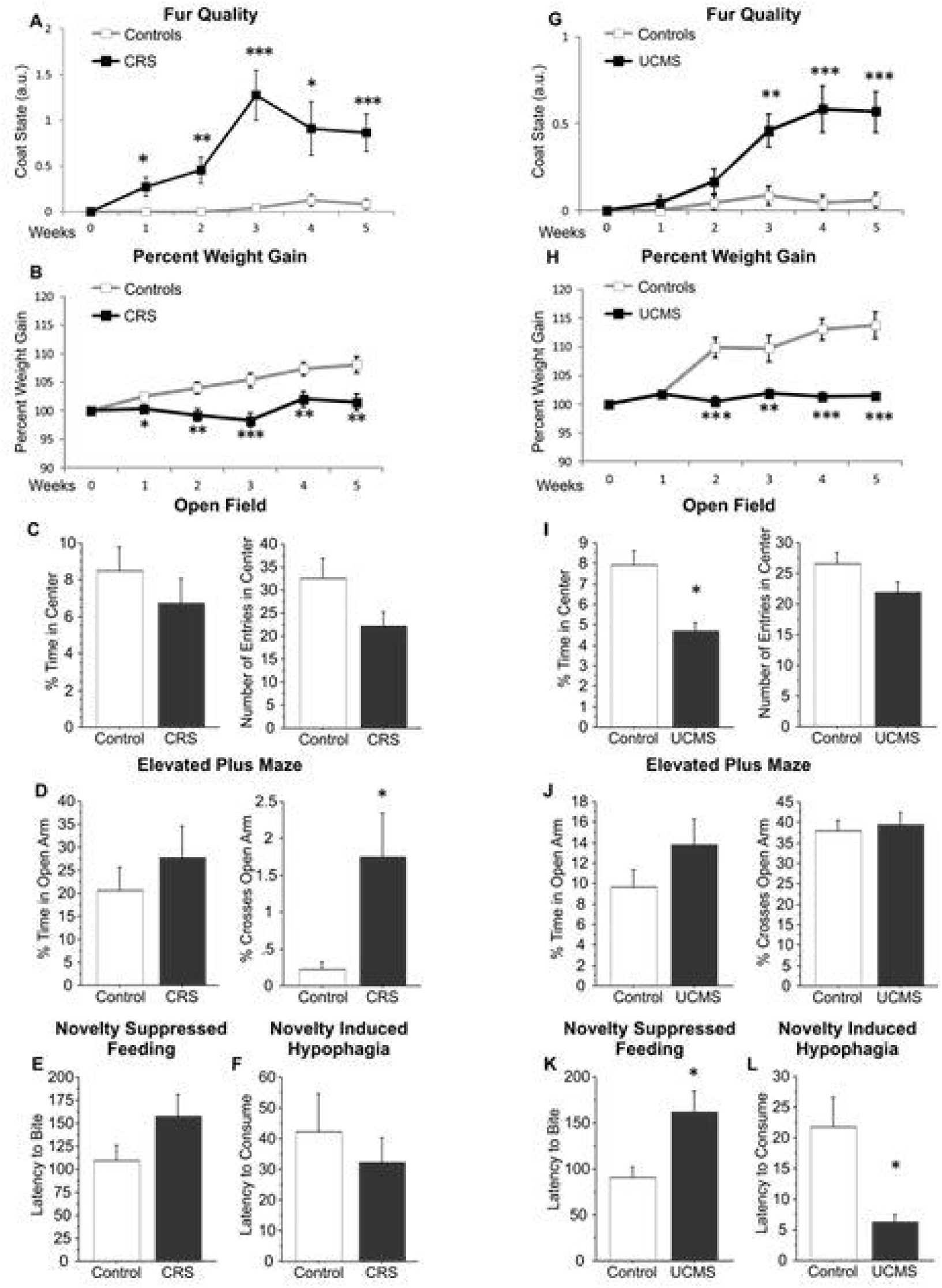
Physical and behavioral changes induced by two chronic stress paradigms. Mice were subjected to chronic restraint stress (CRS; A-F) or unpredictable chronic mild stress (UCMS; G-L). Fur quality (A,G) and weight gain (B,H) were assessed on a weekly basis, throughout the stress exposure. After the 5 weeks of chronic stress exposure, mice were tested in the open-field test (C, I) in which the time and the number of entries in the center were quantified. Mice were also tested in the elevated plus maze (D, J) in which the time and the number of crosses in the open arms were assessed. Mice were also tested in the novelty suppressed feeding test (E, K) and the novelty-induced hypophagia (F, L) in which the latency to bite a food pellet or consume the solution was measured. *p < 0.05, **p < 0.01, ***p < 0.001 compared with controls.

#### 3.1.1. Effects of CRS

Repeated-measures ANCOVA of coat state scores revealed a significant main effect of CRS (F_(1;105)_=15.76; *p*<0.001), time (F_(5;105)_=14.09; *p*<0.0001) and a stress*time interaction (F_(5;105)_=10.5; *p*<0.0001) (**Fig. 1A**). *Post-hoc* analysis identified a significant increase in coat state deterioration induced by CRS exposure from week 1 to week 5 (all p<0.05). A significantly sex difference was found, wherein an exacerbated effect of CRS exposure was detected in males compared to females (F_(1;95)_=97.64; *p*<0.0001) between weeks 2 and 5 *(p<*0.001; **Fig. S2**).

In the same cohort, the analysis of weight gain showed a significant effect of CRS (F_(1;105)_=12.04; *p*<0.01), time (F_(5;105)_=15.61; *p*<0.0001) and a stress*time interaction (F_(5;105)_=7.9; *p*<0.0001). A significant decrease was found in weight gain induced by CRS exposure from weeks 1 to 5 *(p<*0.05, **Fig. 1B**). When sex was considered as a covariate, a significant effect was found, showing exacerbated effects of CRS exposure on weight gain in CRS-exposed males compared to females (F_(1;95)_=18.78; *p*<0.001), and compared to control mice starting weeks 2 to 5 of CRS exposure (*p*<0.05; **Fig.S2**).

In the OF test **(Fig. 1C)**, neither the percentage of time in the center, nor the number of entries in the center (**Fig. 1B**) revealed effects of CRS. In the EPM test (**Fig.1D**), mice exposed to CRS displayed a non-significant change in the percent of time spent and entries in open arms. Lastly, NSF and NIH tests did not detect significant differences in the latency to feed or drink, respectively, in home cage or novel environments (**Fig. 1E-F**). Covariate analysis for sex revealed no main effects or interaction with stress in the EPM and NSF. In the OF test, an interaction was found (F_(1;19)_=5.79; *p*<0.05) wherein females showed greater entries in the center than males at baseline (p<0.05), and decreased entries in the center following CRS (p<0.05). In the NIH test, ANCOVA revealed a significant main effect of sex (F_(1;19)_=4.95; *p*<0.05), wherein *post hoc* analysis identified greater latency to drink in females than males (*p*=0.04; **Fig. S2**).

#### 3.1.2. Effects of UCMS

Repeated measures ANCOVA of coat state degradation revealed a significant main effect of stress (F_(1;110)_=14.134; *p*<0.01), time (F_(5;110)_=15.808; *p*<0.0001) and a stress*time interaction F_(5;110)_=10.7; *p*<0.0001). *Post hoc* analysis identified significantly increased coat state deterioration in UCMS-exposed mice from weeks 3 to 5 (*p*<0.01, **Fig.1G**). Considering sex as a covariate revealed a significant effect of sex (F_(5;100)_=13.9; p<0.001), wherein coat state deterioration was greater in UCMS males compared to females at weeks 2, 4 and 5 (p<0.05; **Fig. S3**). The analysis of weight gain showed a significant main effect of stress (F_(1;110)_=23.8; *p*<0.0001), time (F_(5;110)_=24.5; *p*<0.0001) and a stress*time interaction (F_(5;110)_=20.16; *p*<0.0001). A significantly decreased weight gain was found among UCMS mice compared to control mice, from weeks 2 to 5 (*p*<0.01, **Fig.1H**). Sex was not a significant covariate on this measure.

The same cohort was then tested in the OF test (**Fig. 1I**). Statistical analysis revealed an effect of UCMS exposure, characterized by decreased percent of time spent in the center zone (*p*<0.001) and a trend towards decreased number of entries into this zone (p=0.07). In the EPM time spent in open arms and percentage of open arm crosses was not significantly modified by stress (**Fig. 1J**). In the NSF test UCMS exposure induced a significant increase in latency to feed in the novel arena (p<0.05; **Fig. 1K**). However, UCMS induced a decreased latency to consume milk in the NIH (p<0.01; **Fig. 1L**). Home cage latency to drink or feed was unchanged.

Sex was a not a significant covariate in the OF, EPM and NIH. In the NSF, we found a significant sex*stress interaction (F_(1;18)_=4.6; *p*<0.05). *Post hoc* analysis revealed increased latency to feed in UCMS females compared to UCMS males (*p*=0.02, **Fig. S3**).

### 3.2. Baseline behavior and light challenge response in the PhenoTyper test

The PhenoTyper apparatus was used to monitor food and shelter zones time of male and female mice throughout the dark-cycle. One male mouse showed intermittent missing data during tracking acquisition and was excluded from the study. At baseline, repeated measures ANCOVA revealed a significant sex effect on food zone time (F_1,252_ = 4.19; *p*<0.05). However, *post hoc* analysis revealed significant differences only for the 4^th^ hour of the dark cycle (*p*<0.01, **Fig. 2A**). No significant effect of sex was found for shelter zone time (**Fig. 2B**).

**Figure 2:**
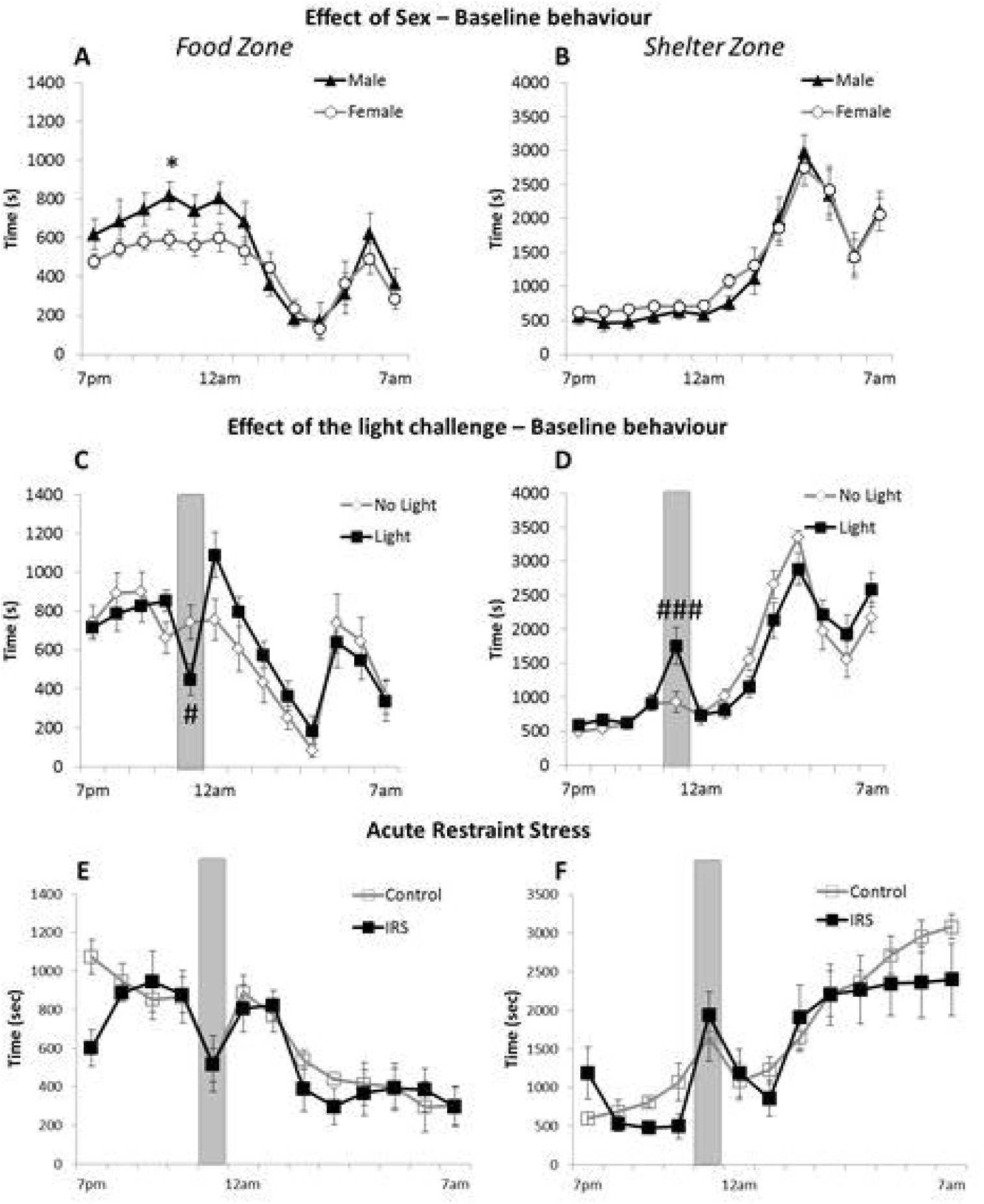
Hourly detection of behavioral changes in the PhenoTyper test. Food zone (A, C, E) and shelter zone (B,D,F) times were assessed every hour. Male and female mice were placed overnight (from 7pm to 7am) in the PhenoTyper boxes where food (A) and shelter (D) zone times are measured. The following week, the same mice were placed in the PhenoTyper boxes, this time with 50% of each sex exposed to a light challenge between 11pm and 12am over the food zone. The same parameters (time in the food (C) and shelter zones (D)) were assessed. Finally, another cohort of mice was subjected to acute restraint stress (ARS) for 1hr, 1hr before being placed in the PhenoTyper boxes (E, F). A light challenge was applied from 11pm to 12am, over the food zone. Time spent in the food and shelter zone was quantified. *p < 0.05 compared to males, # p < 0.05, ### p < 0.001 compared to group receiving no light.

One week after baseline monitoring, mice were placed in the PhenoTyper again and we assessed responses to an acute light challenge applied from 11:00pm-12:00am for half of the animals (*n*=12/group randomly assigned) (**Fig. 2C-D**). The analysis of food zone time revealed a significant time*“light condition” interaction (F_(1;252)_ =1.971; *p*<0.05). The 1h light challenge significantly decreased the food zone time during this time point (*p*<0.05, **Fig. 2C**). ANCOVA also revealed a significant time* “light condition” interaction, wherein the light challenge induced a significant increase in shelter zone time when the light was on, compared to the behavior of animals not experiencing the light challenge (*p*<0.001, **Fig. 2D**). No main effect of sex or sex*stress interaction was found.

### 3.3. Acute restraint stress (ARS) does not alter baseline behavior and light challenge response in the PhenoTyper test

In a separate cohort, we tested the behavioral response to a 1 hour ARS performed immediately before placing the animal in the PhenoTyper apparatus (**Fig. 2E-F**). Analysis of the food (**Fig. 2E**) and shelter time (**Fig. 2F**) did not reveal differences between groups or a time*stress interaction (F_(1;128)_=1.18 and F_(1;168)_=0.94, respectively), suggesting that ARS animals behaved in this test in a similar manner to control mice at baseline, during or following the light challenge. No sex difference was found. Note that ARS induced a decreased food zone time and a mirror increased shelter zone time in ARS-exposed animals from 7 to 8pm, however this effect was not significant since the analysis was performed over the 12hours monitoring.

We also tested whether stress resulting from single housing alters behavior in this test and found no significant difference in shelter or food zone time between control single-housed and control group-housed mice (**Fig. S4**).

### 3.4. Characterization of the trajectory of behavioral changes induced by chronic stress in the PhenoTyper test

#### 3.4.1. Effects of CRS

Repeated measures ANCOVA revealed a significant effect of CRS exposure after 1 week of CRS (**Fig. 3A**; F_(1;252)_ = 9.49; *p*< 0.01) that was consistent for every following week (week 2, F_(1;252)_ = 35.09; *p*<0.0001, **Fig. 3B**; week 3, F_(1;252)_ = 19.9; *p*<0.001, **Fig. 3C**; week 4, F_(1;252)_ = 35.56; *p*<0.0001, **Fig. 3D**; week 5, F_(1;252)_ = 20.92; *p*<0.001,**Fig. 3E**). CRS-exposed mice spent more time in the shelter zone at several time points between 12:00 and 4:00 (following the light challenge) for every week of testing (**Fig. 3**). A similar analysis performed on food zone time revealed stress*time interactions at week 1 (F_(1;252)_=2.29; *p*<0.01), week 2 (F_(1;252_ =1.7; p<0.05), week 3 (F_(1;252)_=2.9; *p*<0.001), week 4 (F_(1;252)_ = 2.6; *p*<0.01) and week 5 (F_(1;252)_ = 2.8; *p*<0.001) (**Fig. S5**). CRS-exposed mice spent less time in the food zone at several time points between 12:00 and 4:00 (following the light challenge) for every week of testing (**Fig. S5**). No main effect of sex or sex*stress interaction were found.

**Figure 3:**
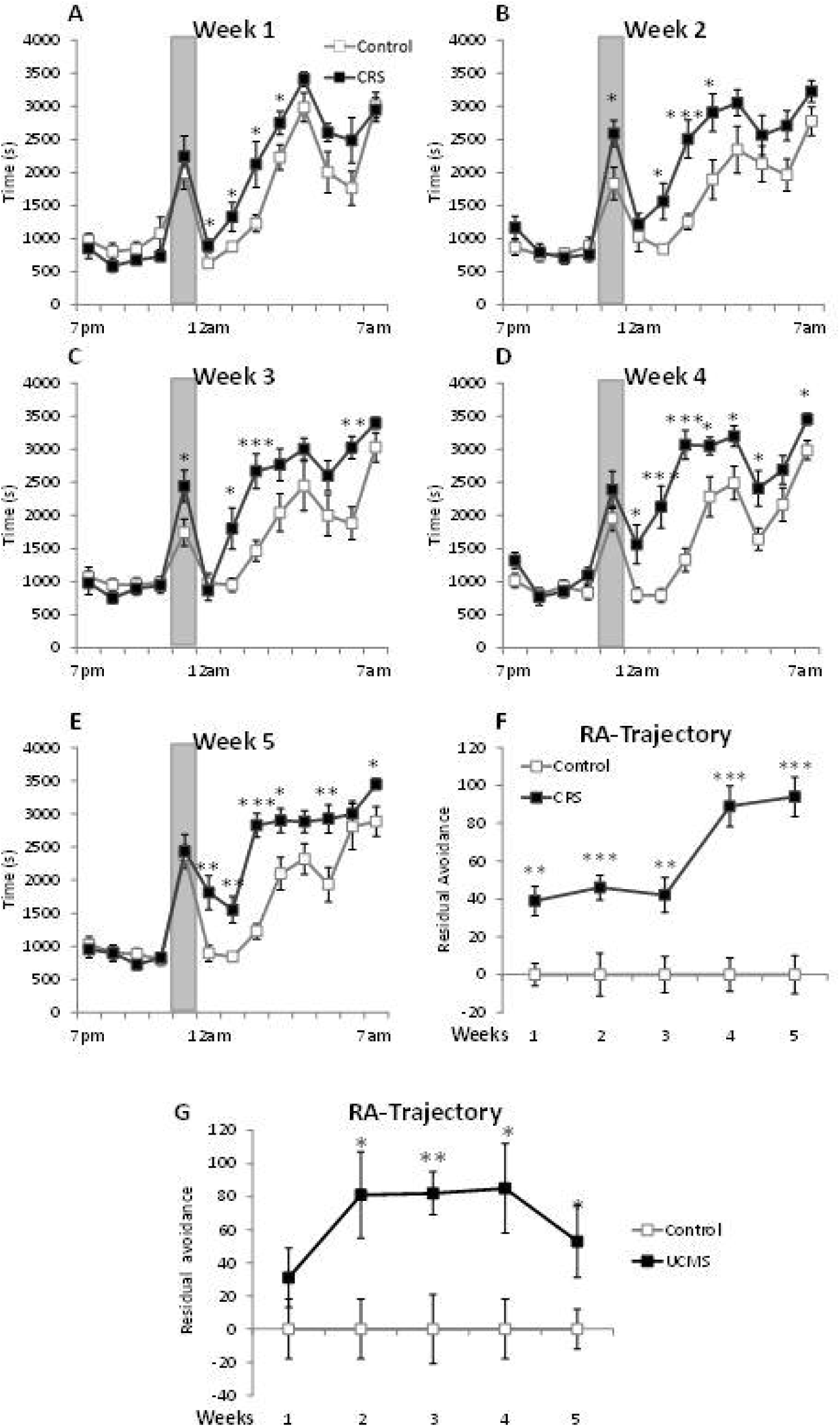
Chronic restraint stress induces behavioral alterations of the shelter zone time in the PhenoTyper test. Mice were subjected to chronic restraint stress (CRS) for 5 weeks, and tested in the PhenoTyper test every week with a light challenge occurring between 11pm and 12am. The shelter zone time was assessed every week (A-E). Using the residual avoidance (RA) calculation, each week’s shelter zone time was transformed into a single value resuming the trajectory of the effect of chronic stress exposure (F). The same calculation was applied to the shelter zone time of mice subjected to unpredictable chronic mild stress (UCMS; G). *p < 0.05, **p < 0.01, ***p < 0.001 as compared with controls.

#### 3.4.2. Effects of UCMS

Repeated measures ANCOVA revealed a significant effect of UCMS exposure after 1 week of UCMS (F_(1;264)_ = 10.172; p<0.0001) that was consistent for every following week (week 2, F_(1;264)_ = 10.790; p<0.0001; week 3, F_(1;264)_ = 16.34; p<0.001; week 4, F_(1;264)_ = 6.569; p<0.05; week 5, F_(1;264)_ = 12.608; p<0.01) in the shelter zone. UMCS-exposed mice spent more time in the shelter zone at several time points following the light challenge (between 12:00 and 4:00) for every week of testing (**Fig. S6**). Mirror results were found in the food zone (i.e. reduced time), where a significant effect of UCMS was observed in week 1 (F_(1;264)_ = 4.624; p<0.05), week 2 (F_(1;264)_ = 11.546; p<0.01), week 3 (F_(1;264)_ = 6.771; p<0.05), week 4 (F_(1;264)_ = 5.833; p<0.05) and week 5 (F_(1;264)_ = 8.413; p<0.01) (**Fig. S7**). No main effect of sex or sex*stress interaction was found.

In a separate cohort, we verified that UCMS-induced reduced time in the food zone was not due to lack of appetite or change in total food consumption. We found no changes in total food intake during the light or dark cycle in home cage, and no changes in overnight food consumption if the animals were placed in the PhenoTyper apparatus without application of the light challenge (**Fig. S8**). However, UCMS mice exposed to the light challenge in the PhenoTyper apparatus showed a significant decreased in overnight total food intake (**Fig.S8**).

#### 3.4.3. Residual avoidance (RA) provides a summary readout of CRS- and UCMS-induced behavioral deficits in the PhenoTyper test

Interestingly, the most striking differences between control and chronic stress-exposed groups were not found before, or during the light challenge, but in the hours following the light challenge. Chronic stress-exposed mice continued to hide in the shelter zone after the challenge and did not return to control group levels until later in the monitoring period. To establish a method for analyzing and presenting these post-challenge differences, we developed the “Residual Avoidance” (RA) calculation. RA illustrates the difference in shelter time post-challenge between chronic stress-exposed and control mice for each week (**Fig. 3F**). Repeated measures ANCOVA revealed a significant effect of stress exposure (F_(1,84)_=49.31; *p*<0.0001). *Post hoc* analysis revealed increased RA among CRS mice for each week (all *p*>.0.001). This effect was highly consistent between and within groups and sexes (**Fig. S9**). Mirror results were obtained when RA calculations were applied to food zone time. Analysis of food zone RA revealed a significant effect of stress exposure (F_(1;84)_=16.5; *p*<0.001). Here, RA was significantly increased after 1, 4, and 5 weeks of CRS (*p*<0.01, **Fig. S5**), with no sex differences on this measure.

Analysis of shelter zone RA in the UCMS model provided similar results, as summarized in **Fig. 3G**. A significant effect of UCMS exposure (F_1;88_=15.5; *p*<0.001) was found without sex or stress*sex interaction. UCMS induced a significant increase in RA at 2 weeks (*p*<0.01) and for every subsequent week of testing (*p*<0.05). It is important to mention that in this cohort, despite a clear group effect, 2 males UCMS out of 12 UCMS animals showed a consistent absence of RA deficits suggesting that RA may be a usable parameter to distinguish susceptible vs. resilient animals (**Fig. S10**). Finally, analysis of food zone RA revealed a significant main effect of stress-exposure (F_1;88_=6.8; *p*<0.05) with no main effect of sex or stress*sex interaction. Analysis identified a significant increase in RA for UCMS-exposed mice at the 4 week time point (*p*<0.05, **Fig. S7**).

### 3.5. Chronic stress exposure, and its interaction with sex, account for the majority of variance across 16 behavioral test parameters

We employed dimension reduction via principal component analysis (PCA) to identify the main factors contributing to the variance among behavioral tests employed in each of the aforementioned experiments, and to determine if PhenoTyper parameters such as RA belong to similar categorical dimensions as parameters measured with commonly-used tests. PCA of 16 behavioral parameters indicated 2 components capturing 25.5% (PC1) and 21.2% (PC2) of behavioral variance across CRS and control groups (**Fig. S11**). PC1 had the strongest loadings (absolute values, >0.4) from anxiety/weight gain-related variables in individual tests and weaker (<0.4) not significant loadings from variables such as fur coat deterioration, home cage latency to feed or drink, or PhenoTyper pre-challenge behavior (**Table 1**). ANCOVA revealed that CRS-exposed mice had significantly higher PC1 scores compared to controls (F_(1;19)_=22.5; *p*<0.0001; **Fig. 4A**) with no main effect of, or interaction with, sex (**Fig. S11**). PC2 captured variance from the majority of the commonly-used tests employed as well as that from home cage feeding or drinking, and fur coat deterioration. ANCOVA of PC2 scores revealed significant main effects of stress (F_(1;19)_=12.4; *p*=0.002), sex (F_(1;19)_=10.0; *p*=0.005) and a stress*sex interaction (F_(1;19)_=10.0; *p*<0.01, **Fig. 4A and Fig. S11**), wherein CRS males had significantly lower PC2 scores than CRS females, but there were no sex differences among control animals. Our results demonstrated that animals with higher PC1 scores had greater elevations in emotional reactivity across dimensions and groups, whereas PC2 scores differentiated the influence of sex for CRS groups (**Fig. 4A**).

**Figure 4:**
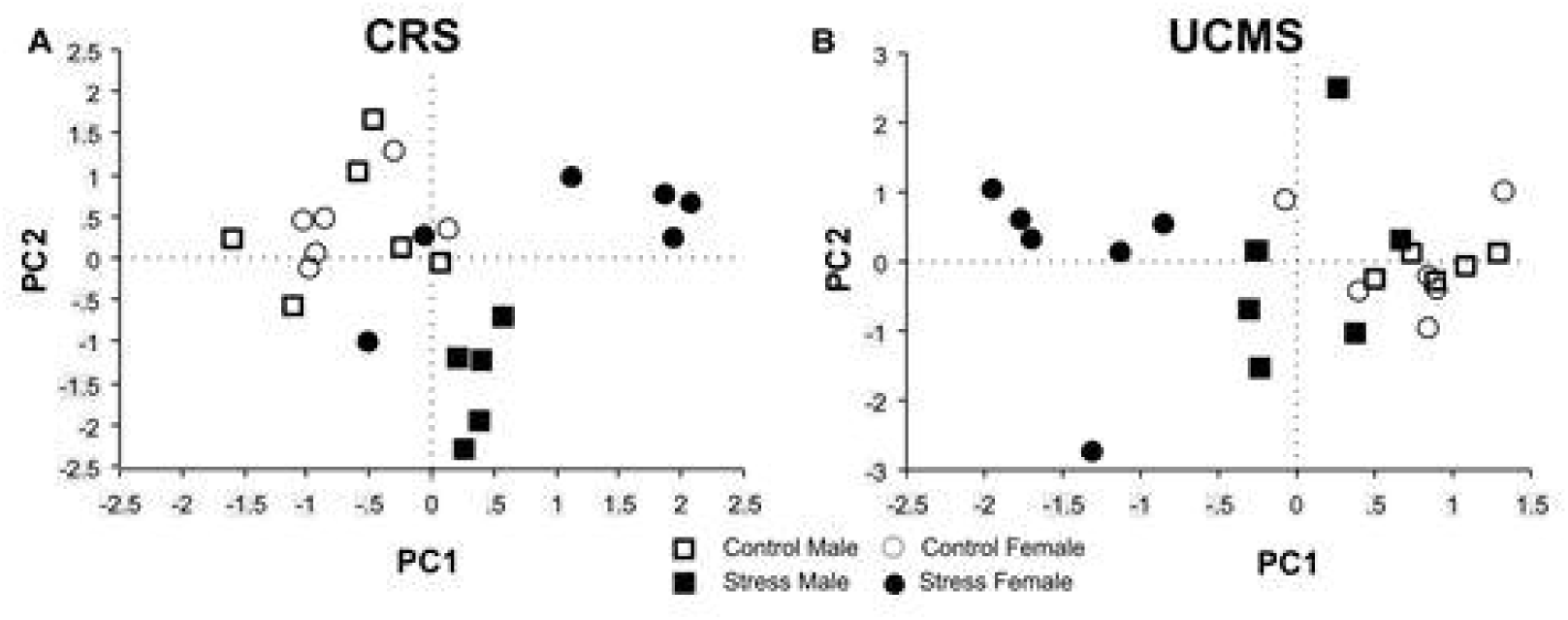
Principal component (PC) analysis of behavioral data obtained from mice exposed or not to chronic stress. Data obtained from animals subjected to chronic restraint stress (CRS) and respective control group tested in the elevated plus maze, open-field, novelty suppressed feeding, novelty-induced hypophagia and the PhenoTyper test were used for factor analysis (A). PC1 and PC2 captured behavioral variance of stress and sex across groups. Similar factor analysis was performed on data collected from mice subjected to unpredictable chronic mild stress (UCMS) and respective control group. PC1 captured the behavioral variance of stress across groups. (B).

PCA was also performed on data collected from the UCMS cohort. In this case, PCA of 16 variables indicated 2 components capturing 27.3% (PC1) and 17.4% (PC2) of behavioral variance across UCMS and control groups (**Fig. S11**). ANCOVA of PC1 scores revealed significant main effects of stress (F_(1;20)_=53.1; *p*<0.0001), sex (F_(1;20)_=16.1; *p*<0.001) and a stress*sex interaction F_(1;19)_=16.1; *p*<0.001, **Fig. 4B and Fig. S11**). No effects of stress, sex, or and stress*sex interaction were found for PC2 scores (**Fig. S11)**. Our results indicate that animals with lower PC1 scores had greater emotional reactivity across dimensions and groups, wherein UCMS female mice displayed lower PC1 scores than UCMS males and control animals (**Fig. 4B**). In both experiments there were other components with eigenvalue >1, which were, however, independent of sex and/or stress (**Fig. S11**) and probably capture variance from other sources.

Finally, the loadings of the variables derived from the commonly-used tests were distributed on the main 2 PCs of each experiment (**Table 1**); therefore it is difficult to determine the precise behavioral significance of each individual PC. Yet, it is interesting to note that only shelter zone RA and percent time in the center in the OF specifically loaded on the stress dependent PCs in both cohorts (**Table 1**). Indeed, in both studies shelter zone RA loaded strongly on the PCs that captured the behavioral variance attributed to chronic stress (i.e. PC1 and PC2 for the CRS experiment and PC1 for UCMS experiment), along with the variables that measured approach/avoidance conflict or aversion in the commonly-used tests. These results suggest that the shelter RA may belong to similar categorical dimensions as behavioral parameters measured with commonly-used tests known to be affected by chronic stress.

### 3.6. CRS-induced increases in residual avoidance are reversed by treatment with chronic imipramine, but not acute diazepam

A separate cohort of mice was added to identify the effects of DZP in control and CRS conditions (**Fig. 5A-B**). Analysis of shelter zone time revealed a significant main effect of stress (F_(1;504)_=12.76; *p*<0.001), no main effect of drug treatment (F_(1;504)_=0.15; *p*=0.69), and an interaction between stress*drug conditions (F_(1;504)_=7.5; *p*<0.01). However, *post hoc* analysis revealed that CRS mice spent significantly more time in the shelter zone regardless of drug treatment (p<0.05). Despite a significant stress*drug interaction, no significant differences were identified at particular time points between control or CRS mouse group receiving vehicle or DZP. However, RA analysis revealed a significant main effect of stress (F_(1;42)_=26.29; *p*<0.0001) and its interaction with drug treatment (F_(1;42)_=4.3; *p*<0.05), wherein *post hoc* analysis showed decreased RA in control animals receiving DZP (*p*<0.05, **Fig. 5A**). We further confirmed increased RA among CRS mice (*p*<0.01) and found no significant differences between CRS animals treated with vehicle or DZP. No significant effect of sex was found. It is important to mention that the lack of effects of DZP does not appear to be dependent on the dose or timing of administration. Indeed, in the separate cohort we found that at 3 mg/kg, i.p., DZP again failed to reverse CRS-induced behavioral alterations (data not shown). We also assessed the effects of DZP (1.5 mg/kg, i.p) 30mins before the light challenge and found no effects of this regimen in CRS animals (data not shown).To determine if DZP could reverse the effects of CRS in another test, mice were tested in the NSF (**Fig. 5C**). ANCOVA revealed significant main effects of stress (F_(1;48)_=24.7; *p*<0.001) and drug treatment (F_(1;48)_=7,1; *p*<0.05), but no stress*drug interaction. In this experiment, CRS significantly increased the latency to feed (p<0.05). In addition, DZP had no effects on control animals, but significantly reduced the latency to feed in CRS animals (*p*<0.05). Home cage latency to drink or feed was unchanged. Consideration of sex as covariate revealed a main effect of stress (F_(1;44)_=13.6; *p*<0.001) and a stress*sex interaction (F_(1;44)_=3.3; *p*<0.001), but no effect of drug*stress*sex interaction for latency to feed, reflecting a greater increase in latency to feed among CRS females compared to males.

**Figure 5:**
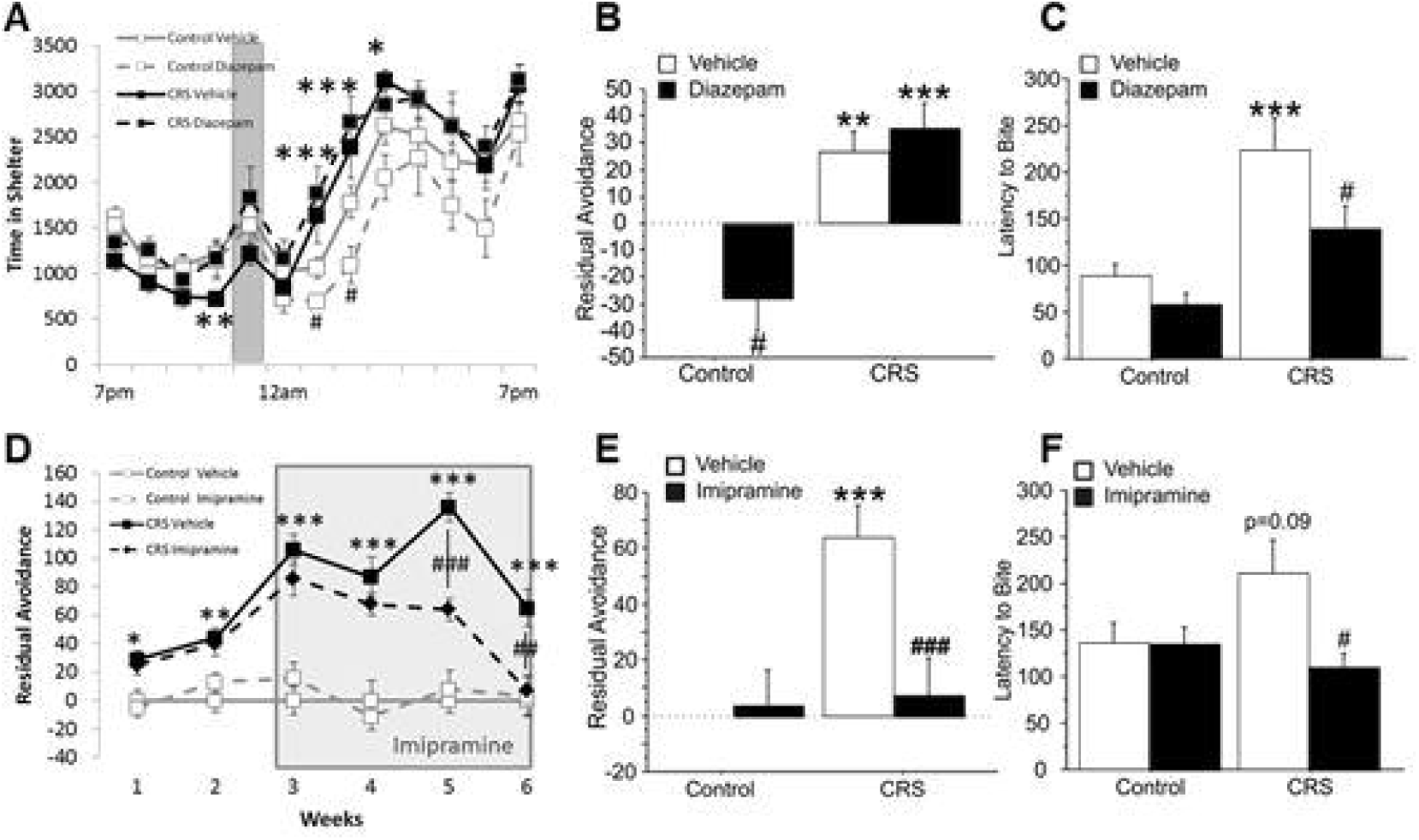
Reversal of behavioral deficits after anxiolytic or antidepressant treatments. Mice were tested in the PhenoTyper test (A-B) and in the novelty-suppressed feeding (C) after being subjected to chronic restraint stress (CRS). Animals also received acute diazepam treatment 30min prior being placed in the apparatus. The shelter zone time was quantified in the PhenoTyper test (A), and the residual avoidance was calculated based on these results obtained in the shelter zone (B). Mice were also tested in the novelty-suppressed feeding, in which the latency to bite was monitored. A separate cohort of animals was subjected to CRS and received imipramine chronically in the drinking water after the second week on CRS exposure, and for the next 4 weeks. Residual avoidance in the shelter zone was calculated for every week (D). Effect of imipramine on shelter RA obtained after 4 weeks of treatment was quantified (E). Finally, after 6 weeks of CRS and 4 weeks of imipramine treatment, mice were tested in the novelty-suppressed feeding test in which the latency to bite the food pellet was measured (F). *p < 0.05, **p < 0.01, ***p < 0.001 compared to control animals #p < 0.05, ##p < 0.01, ###p < 0.001 compared to vehicle treated animals.

In an additional cohort, we replicated the effects of CRS on RA throughout 6 weeks of stress exposure (F_(1;220)_=72.7; *p*<0.0001), and assessed the effects of chronic imipramine treatment starting on day 21. We found a significant stress*drug (F_(1;220)_=5.4; *p*=0.023) and stress*drug*time interactions (F_(5;220)_=3.25; *p*=0.007) on RA. There was no significant effects of imipramine at week 3 (after 1 day of treatment) or at week 4 (after 1 week of treatment) in control or CRS mice (*p*>0.05, **Fig. 5D**). However, a significant difference between vehicle- and imipramine-treated CRS mice was identified on weeks 5 and 6 (after 2 and 3 weeks of treatment, respectively; *p*<0.01, **Fig. 5D and E**). No significant effect of sex was found.

To determine if imipramine could reverse CRS effects in another test, mice were then tested in the NSF test. ANCOVA revealed a significant main effect of drug (F_(1;44)_=4.5; *p*<0.05), and a stress*drug interaction (F_(1;44)_=4.4; *p*<0.05). In this experiment, CRS induced a trend toward an increased latency to feed (p=0.09). In addition, chronic imipramine had no effect in non-stress animals, but significantly reduced the latency to feed in CRS animals (*p*<0.05, **Fig.5F**). Home cage latency to drink or feed was unchanged. Analysis of sex as a covariate revealed a significant main effect (F_(1;44)_=14.7; *p*<0.001), a stress*drug interaction (F_(1;44)_=7.8; *p*<0.01), and a drug*stress*sex interaction (F_(1;44)_=8.8; *p*<0.01) on latency to feed. These changes were driven by an overall greater increase in latency to feed following CRS and decreased latency following imipramine in females compared to males.

## 4. Discussion

Our findings confirmed that mice subjected to UCMS or CRS displayed clear responses to chronic stress exposure on longitudinal physical readouts such as weight gain and coat state deterioration. However, when tested under similar experimental conditions, common tests used to assess chronic stress-induced behavioral deficits such as the EPM, OF, NSF, and NIH showed highly heterogeneous results between tests and stress models. We also assessed the effects of chronic stress in a recently developed test allowing for monitoring of the animal’s normal exploratory behavior before, during and after a light challenge in a home cage-like setting, called the PhenoTyper test (based on many other tests named after their cognate apparatus). In this test, mice subjected to UCMS or CRS showed a common response after the light challenge, i.e. an avoidance of the lit zone in favor of a shelter zone, lasting for hours beyond termination of the challenge. Notably, this “residual avoidance” (RA) after the light challenge was replicated across the 4 experiments performed in this study using 2 stress models, including in both males and females. Stress-induced elevations in RA were detected rapidly, following the first/second week of UCMS or CRS exposure, and importantly were consistently detected in the same animals when tested a week apart for up to 5/6 weeks. RA was not altered by acute restraint stress, suggesting the need for biological effects of chronic stress. Acute DZP reduced RA in control but not chronic stress conditions. On the other hand, chronic treatment with imipramine (2-3 weeks), but not acute treatment with DZP or imipramine, was sufficient to reverse CRS-induced RA. Altogether, our findings demonstrate that shelter RA is a highly consistent and repeatable readout for chronic stress-induced “emotional reactivity” that responds to chronic antidepressant treatment.

The main objectives of the current study were: 1) to fully characterize a recently developed experimenter-free, repeatable, behavioral test that consistently detect the effects of chronic stress, 2) to compare chronic stress-induced behavioral changes among the most widely-used tests with those measured by the PhenoTyper test, 3) to determine whether the PhenoTyper test could identify similar profiles of elevated emotional reactivity between two well-documented chronic stress models and, 4) to investigate the effects of an anxiolytic and an antidepressant in this test.

In designing experiments, researchers generally choose a chronic stress model and one (or a few) test(s) that produces consistent results under lab-specific conditions. This approach has proven its usefulness, since previous work reports increased anxiety/depressive-like deficits following chronic stress, including studies from our lab (Banasr et al., 2010; Duric et al., 2017; Nikolova et al., 2018; Soumier and Sibille, 2014). However, here we demonstrate that UCMS or CRS procedures, while clearly effective (i.e. robust effects on coat state and weight gain assessments) induced highly variable effects on anxiety-like behaviors across multiple behavioral tests, even after 5 weeks of stress exposure. In accordance with published and unpublished data around the world, we confirmed inconsistent chronic stress effects in those acute tests, arguing against (or in favor of, depending on the reasoning) the idea that having multiple tests assessing the same phenotype is the best strategy (Ramos, 2008). The lack of reliable effects of chronic stress in these tests is often blamed on issues with chronic stress models, stress strength or length, mouse strain, and/or sex differences or how the tests were performed (Castanheira et al., 2018; Cryan and Sweeney, 2011; Ferreira et al., 2018; Willner, 2017a, b).To compensate for this potential bias, scientists have developed several other behavioral tests/measurements which capture other dimensions of the chronic stress response including anhedonia (Willner, 2005; Mineur et al 2006; Surget et al. 2011, Mateus-Pinheiro et al 2014)

Standard tools that measure anxiety-like behavior provide a snapshot of the animal’s behavior (Ohl, 2005), and are highly sensitive to experimenter-bias with regards to how animals are handled before and during testing. Indeed, given that each previous time a mouse from the stress group was handled a stressor was applied, animals may naturally develop a rapid and transitory experimenter-induced hyperlocomotor activity. Since most commonly-used tests are short and highly dependent on exploratory drive, more often than not animals subjected to CRS and UCMS may display a greater number of entries in open arms of the EPM or in center zone of the OF, results usually interpreted as decreased anxiety-like behavior. In addition, it is not uncommon to find (as illustrated in this study) that the same UCMS animals display opposite phenotypes in very similar tests such as the NSF and NIH or the EPM and OF. In these cases, drawing conclusions about the impact of chronic stress from these tests can be difficult. Since chronic stress-induced behavioral effects are expressed heterogeneously, recent studies have employed z-emotionality scores (averaging z-normalized scores across multiple tests) to measure consistency (or not) of behavioural responses across tests and to establish a type of quantitative scale for emotional reactivity to stress (Guilloux et al., 2012); loosely comparable to the Hamilton anxiety or depression scales used in humans (DSM-V, 2013).

Here, we used the PhenoTyper test based on the rationale that a longer automatized test in a home-cage-like setting would address the aforementioned caveats and provide a more reliable readout of chronic stress-induced behavioral deficits. This home cage-like setting also offers the unique advantage of being non-reliant on novelty, therefore allowing repetitive testing. This test shares several features with light/dark box tests (e.g., using light as stressor to measure avoidance behavior), with the added benefit of monitoring animal behavior for a relatively extended period of time (overnight), and before, during and after the stressor experience (the light challenge). We previously used the PhenoTyper test to assess UCMS effects longitudinally in BalbC male mice (Nikolova et al., 2018) and as a single readout in male and female C57B6 mice (Maluach et al., 2017). In BalbC mice, we found increased shelter zone time before, during, and after the light challenge (Nikolova et al., 2018) and in C57B6 mice, a reduction in shelter zone time after the light (Maluach et al., 2017). Here, the weekly monitoring of C57B6 mice revealed no major effects of either chronic stress model before the light challenge, and a variable increased response during the challenge exhibited some weeks and not others. We believe that this non-systematic effect of CRS or UCMS during the challenge is akin to findings one may see in commonly-used tests, and is likely due to the inherent difficulty of reliably capturing differences during a short testing period. It is also possible that some animals might be more susceptible to acute exposure to stress than others; however, in depth analysis of the weekly testing was not able to identify CRS or UCMS mice showing clear stress resiliency during the light challenge (data not shown). Notably, using RA as readout we could identify mice consistently displaying resiliency to UCMS (2/12). Further testing would be required to determine if a longer or a shorter light challenge may modulate the consistency of the findings for both UCMS and CRS models during the light challenge.

Importantly, UCMS and CRS induced consistent effects on RA in C57B6 mice. This finding was replicated using the UCMS model in parallel experiments with BalbC mice (data not shown), and by analysing the RA of animals from our previous two studies (Maluach et al., 2017; Nikolova et al., 2018). The fact that this deficit was found using both models and both strains and lasted throughout stress exposure, but was not observed after an acute stressor, suggests that it is highly specific to chronic stress. The underlying cause of this maladaptive inability to resume normal behavior is open to speculation. It is clear that a rapid response to an aversive stimulus helps organisms avoid potential threat or harm, but the ability to flexibly adapt to changing environments after the stress response is crucial for the capacity to recover from a challenge or stressor (Feder et al., 2009; McEwen et al., 2015). In this context, RA could be a readout for excessive stress generalisation, which when maladaptive can lead to fear responses that are too strong or occur in inappropriate situations. Indeed, chronic stress exposure was shown to enhance fear learning, impair extinction or induce exacerbated fear responses even in novel contexts or when subjected to heterotypic challenge (Macht and Reagan, 2018; Hoffman et al., 2014, Valentine et al., 2008). Inappropriate fear is a cardinal feature of anxiety disorders (Dunsmoor and Paz, 2015; Lissek, 2012; Luyten et al., 2011), and found in other stress-related illnesses, such as post-traumatic stress and major depressive disorders (DSM-V, 2013). One could also interpret increased RA as enhanced freezing, decreased exploration, and disengagement of the animal towards its environment which would seem related to human neuropsychiatric symptoms such as psychomotor retardation or loss of interest in usual activities, such as food consumption or exploration (i.e., anhedonia). However, this is unlikely since we found no changes in home-cage food consumption during the light or dark cycle in this study, nor correlations between recalculated RA and sucrose consumption measured on animals from our previous studies (Nikolova et al 2018; Maluach et al 2017). Knowing that chronic stress exposure can alter feeding profiles without changing total food intake (Aslani et al 2015), analysis of food consumption pattern in and out of the PhenoTyper apparatus would be needed to clearly assert this interpretation. Such analysis would also have to be done with and without the light challenge since we found that the challenge reduced food intake as a result of increased RA in UCMS animals.

Finally, RA deficits may involve a learning component where CRS and UCMS animals associate the light challenge (i.e. a light phase within the dark cycle) with the likelihood of being subjected to stressors. This is possible for CRS animals since restraint stress sessions were performed only during the subjective day, but unlikely for the UCMS animals since stressors also occurred during the night. In addition, association learning would also translate into avoidance before the challenge (anticipation), greater magnitude of effects during the challenge every week of testing even in the control group (acquisition) or decreased magnitude of chronic stress-induced effects on RA (habituation); features we did not observe in the present study. Still, whether due to generalized fear, to motivation for food or exploration following an aversive stimulus, or to some type of fear learning association, RA deficits are highly specific to chronic stress exposure and segregate with other types of chronic stress-induced conflict anxiety measures in the PC analysis.

The molecular/cellular substrates of chronic stress-induced RA are unknown. Previous work correlated chronic stress-induced cellular changes in corticolimbic regions such as increased amygdala volume (Nikolova et al., 2018) or cortical DNA/RNA oxidation (Malauch et al, 2017) with behavioral deficits that included altered performances in PhenoTyper test. Chronic stress-induced RA may involve molecular mechanisms associated with increased amygdala activity and cortical cellular damage, such as chronically elevated corticosterone levels (Duvarci and Pare 2007, Jorgensen et al 2017). We believe that acute increase in corticosterone levels is involved in the animal’s response during the light challenge. Indeed, light stimuli applied during the dark cycle were shown to transitory induce increased corticosterone levels which last ~1hr (Mohawk et al, 2007). Increased corticosterone may explain changes in the avoidance behavior during the light challenge. Lasting or chronic elevation of corticosterone levels and dysfunction of the HPA associated with chronic stress (Prevot et al. 2018) may also be responsible of RA deficits. If corticosterone levels influence chronic stress-induced RA magnitude of response, testing animals at a different time of the day may increase variability or yield different results. Finally, chronic stress alters circadian rhythms (Landgraf et al, 2016) which may be an additional dimension that may alter chronic stress-induced effects on this measure. These considerations would be relevant aspects to pursue in future studies.

DZP and imipramine were used to test the predictive validity of RA. DZP reduced shelter RA in control conditions, suggesting that RA contains an anxiety-like component. Consistent with this possibility, the principal component analysis further suggests the existence of i) an anxiety-related behavioral dimension shared with other stress sensitive-parameters that measured approach/avoidance conflict or ii) at the very least, a distinct but still related stress-dependent dimension. Interestingly, DZP had no effect on CRS-induced RA, but was efficient in the NSF test. Although this may appear to contradict the interpretation of RA as readout of anxiety-like behavior, it is important to mention that, in stress-exposed animals, differential effects of acute DZP can be similarly observed in other tests, including in tests as similar as EPM and the elevated T-maze (Lapmanee et al., 2013). DZP also showed lower efficacy in light/dark-related tests while showing clear anxiolytic-like effects in the EPM when tested in the same animals at the same dose (Kshama et al., 1990; Rodgers and Johnson, 1995; Rodgers and Shepherd, 1993; Santucci et al., 1994). One could also suggest that differential DZP effects in the NSF and PhenoTyper tests may reflect different anxiety-like states; one that responds acutely to DZP and the other which may require longer treatment with DZP or that responds to a different class of drugs. Reversal of chronic stress-induced RA by long-term imipramine treatment supports the later hypothesis. Indeed, imipramine is often described as a better option than benzodiazepines for the treatment of generalized anxiety in clinical settings (Huh et al., 2011). It is also possible that RA encompasses a stress-related dimension independent of anxiety such as loss of interest, helplessness, or anhedonia, which are classically more sensitive to antidepressant treatment (Frisbee et al., 2015; Valentine et al., 2008; Willner, 2005). This would explain why treatment with imipramine, but not DZP, reversed chronic stress-induced RA. However, further experiments using other classes and regiments of antidepressants or anxiolytics would be needed to validate either hypothesis.

Finally, while we included sex as a biological variable in all experiments, our studies were not designed to examine sex differences and may be underpowered to identify meaningful changes. We confirmed that weight gain and coat state are primary readouts differentially affected by sex in response to stress (Moench and Wellman, 2017; Nollet et al., 2013; Piantadosi et al., 2016; Stanley et al., 2014). Yet, overall, our results show similar effects in males and females on emotional reactivity following CRS or UCMS. The stress and sex interaction found in the PCA mainly illustrates a greater variability in the response to chronic stress in females vs. males. Greater variability in females is frequently documented and can be attributed to factors such as oestrus cycle, level of stress susceptibility, or the dimension being measured by each test (e.g., consumption, exploratory, conflict, or social) (Diaz et al., 2016; Lam et al., 2018; Seney and Sibille, 2014).

Given the necessity and usefulness of rodent behavioral models, it is evident that better behavioral readouts are needed to detect changes to animals’ regular behavior. Here, we developed and validated a readout that detects replicable and repeatable avoidance deficits after a light challenge following chronic stress exposure. Although the equipment is highly specific to mice, similar apparatus (locomotor activity or open field boxes) could be fitted with spotlight and shelter and adapted to measure residual avoidance in rats or other experimental species. Nevertheless, this behavioral index offers the opportunity to assess anxiety-like behavior longitudinally and can be instrumental to pinpoint the chain of events leading to expression of behavioral deficits associated with chronic stress and/or chronic anxiety in mice, and to potentially screen novel anxiolytic and antidepressant treatments.

## Author Contributions and Acknowledgements

TP, KM, ES and MB designed the study and wrote the manuscript. TP, KM ran the behavioral experiments with help from CF, DC for performing stressors and DN for behavioral analysis. CF and YN contributed to the analysis and interpretation of the results. TP and CF are supported by CAMH discovery fund fellowships. KM was supported by a Centre for Collaborative Drug Research (CCDR)—(University of Toronto) Student Award. DN and CF received Ontario Graduate Scholarships during the studies. MB is supported by a NARSAD young investigator award from the Brain & Behavior Research Foundation and the Campbell Family Mental Health Institute of CAMH. ES received the Brain & Behavior Research Foundation and Canadian Institutes for Health Research (CIHR). YN is supported by a NARSAD Young Investigator Award from the Brain & Behavior Research Foundation and a Koerner New Scientist Award. The project was supported by the Campbell Family Mental Health Research Institute.

