## Supplementary for "Residual avoidance: a new consistent and repeatable readout of chronic stress-induced conflict anxiety reversible by antidepressant treatment"

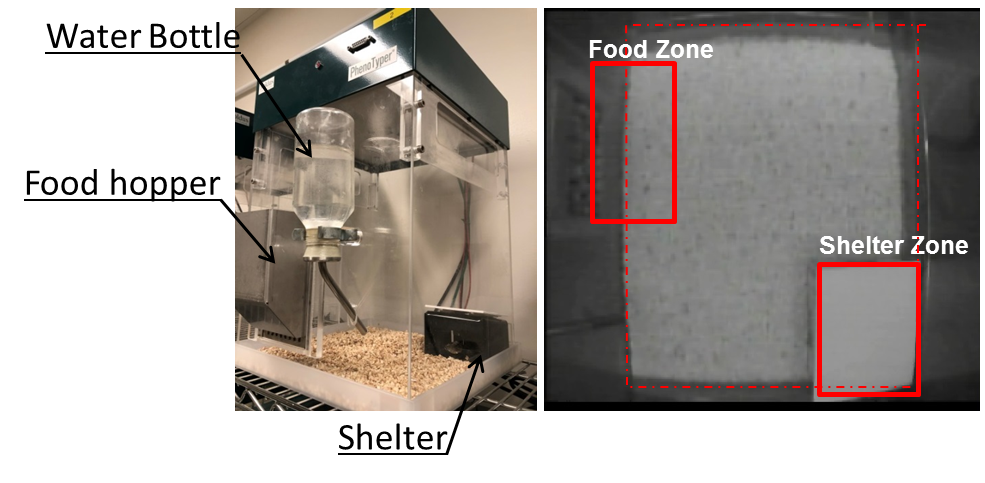

**Figure S1: The PhenoTyper apparatus and delineated zones**

Left panel shows a representative picture of the apparatus taken from the side showing placement of the shelter, the food hopper and the water bottle. The right panel shows a picture of the apparatus taken from above with the delineated zones.

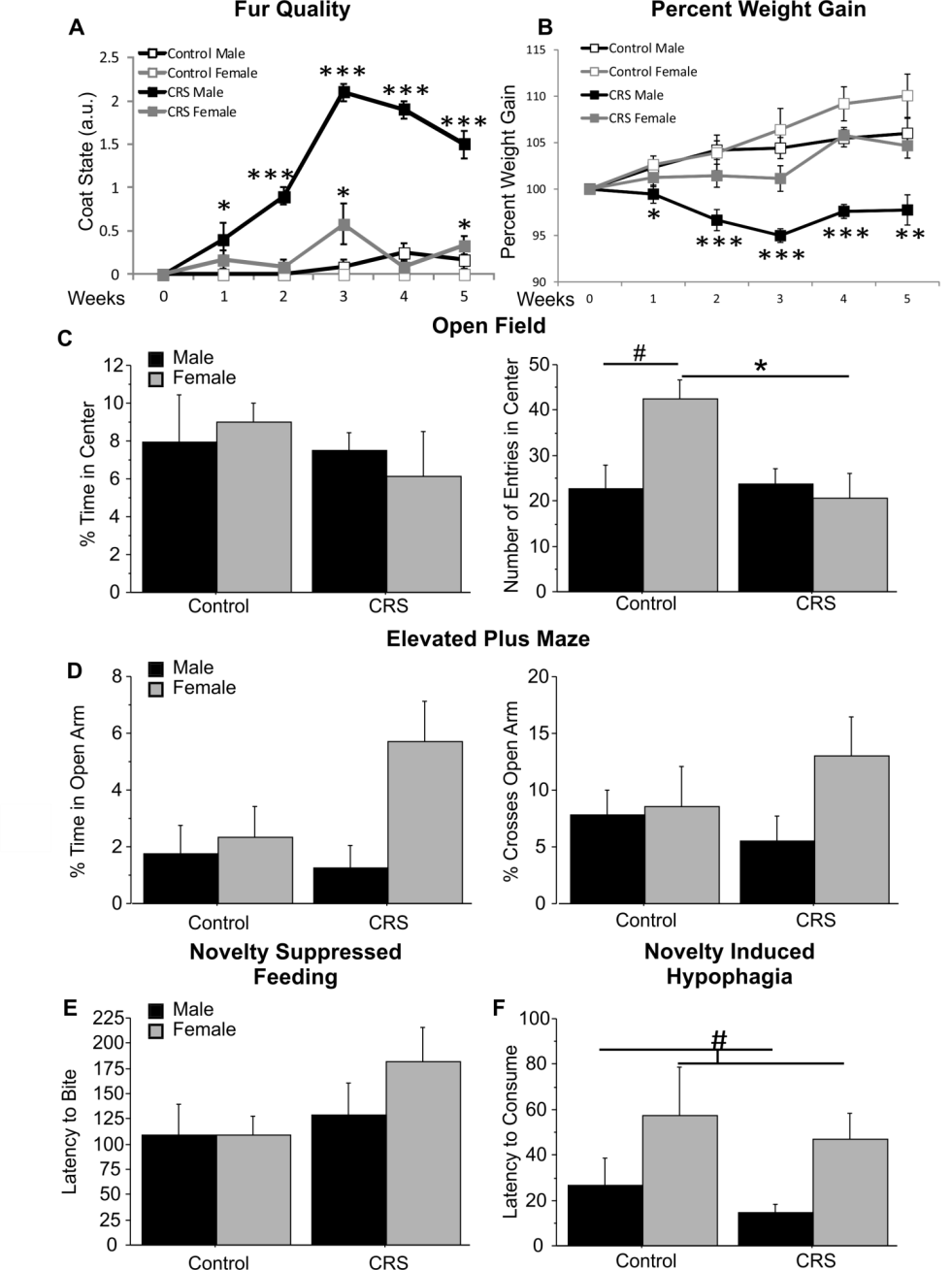

**Figure S2:** **Physical and behavioral changes induced by CRS in males and females.**

Male and female mice were subjected to chronic restraint stress (CRS). Fur quality (A) and weight gain (B) were assessed on a weekly basis, throughout the stress exposure. After 5 weeks of chronic stress exposure, mice were tested in the open-field test (C) in which the time and the number of entries in the center were quantified. Mice were also tested in the elevated plus maze (D) in which the time and the number of crosses in the open arms were assessed. Mice were also tested in the novelty suppressed feeding test (E) and the novelty-induced hypophagia (F) in which the latency to bite a food pellet or consume the solution was measured. *p < 0.05, **p < 0.01, ***p < 0.001 compared with controls. # p < 0.05 compared to males.

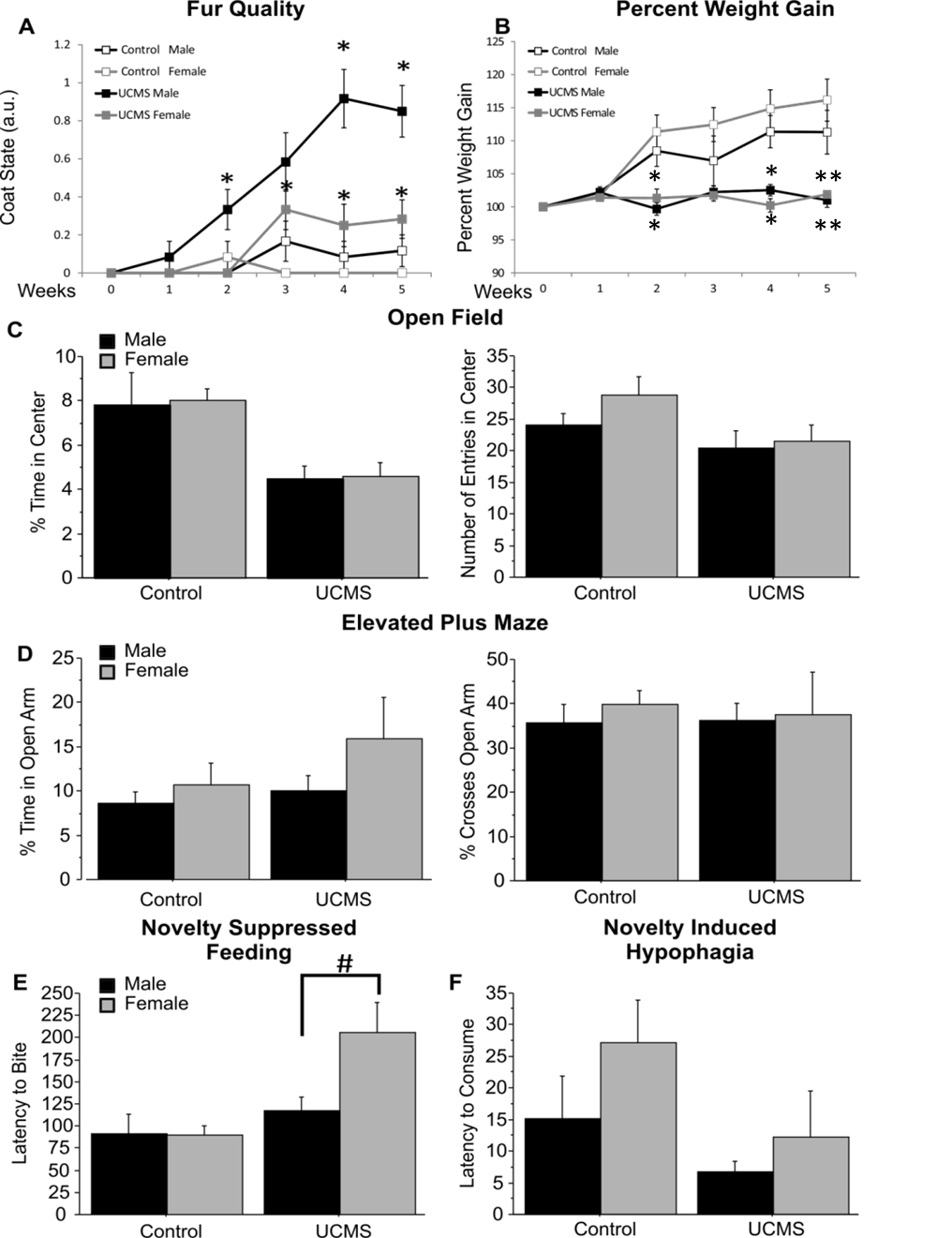

***Figure S3:* Physical and behavioral changes induced by UCMS in males and females.**

Male and female mice were subjected to unpredictable chronic mild stress (UMCS). Fur quality (A) and weight gain (B) were assessed on a weekly basis, throughout the stress exposure. After 5 weeks of chronic stress exposure, mice were tested in the open-field test (C) in which the time and the number of entries in the center were quantified. Mice were also tested in the elevated plus maze (D) in which the time and the number of crosses in the open arms were assessed. Mice were also tested in the novelty suppressed feeding test (E) and the novelty-induced hypophagia (F) in which the latency to bite a food pellet or consume the solution was measured. *p < 0.05, **p < 0.01, ***p < 0.001 compared with controls. # p < 0.05 compared to males.

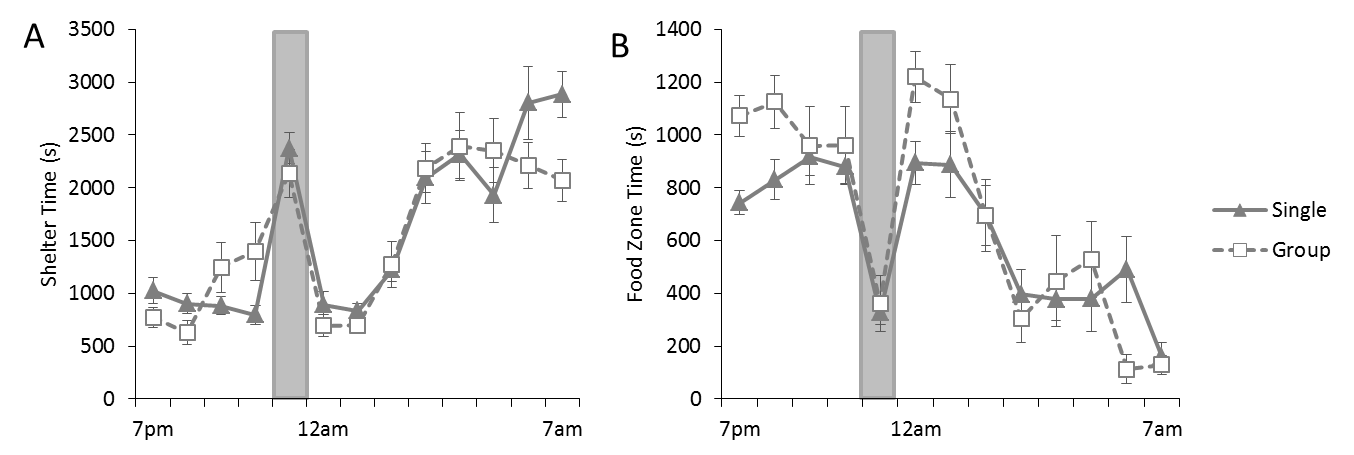

***Figure S4:*** **Single- or group-housing does not alter behavior of control mice in the PhenoTyper test.**

Single housed and group-house mice (3/cage) were tested in the PhenoTyper test (n=12 per group). The time in the Food Zone (A) and the Shelter (B) was measured. No statistically significant difference was found between groups.

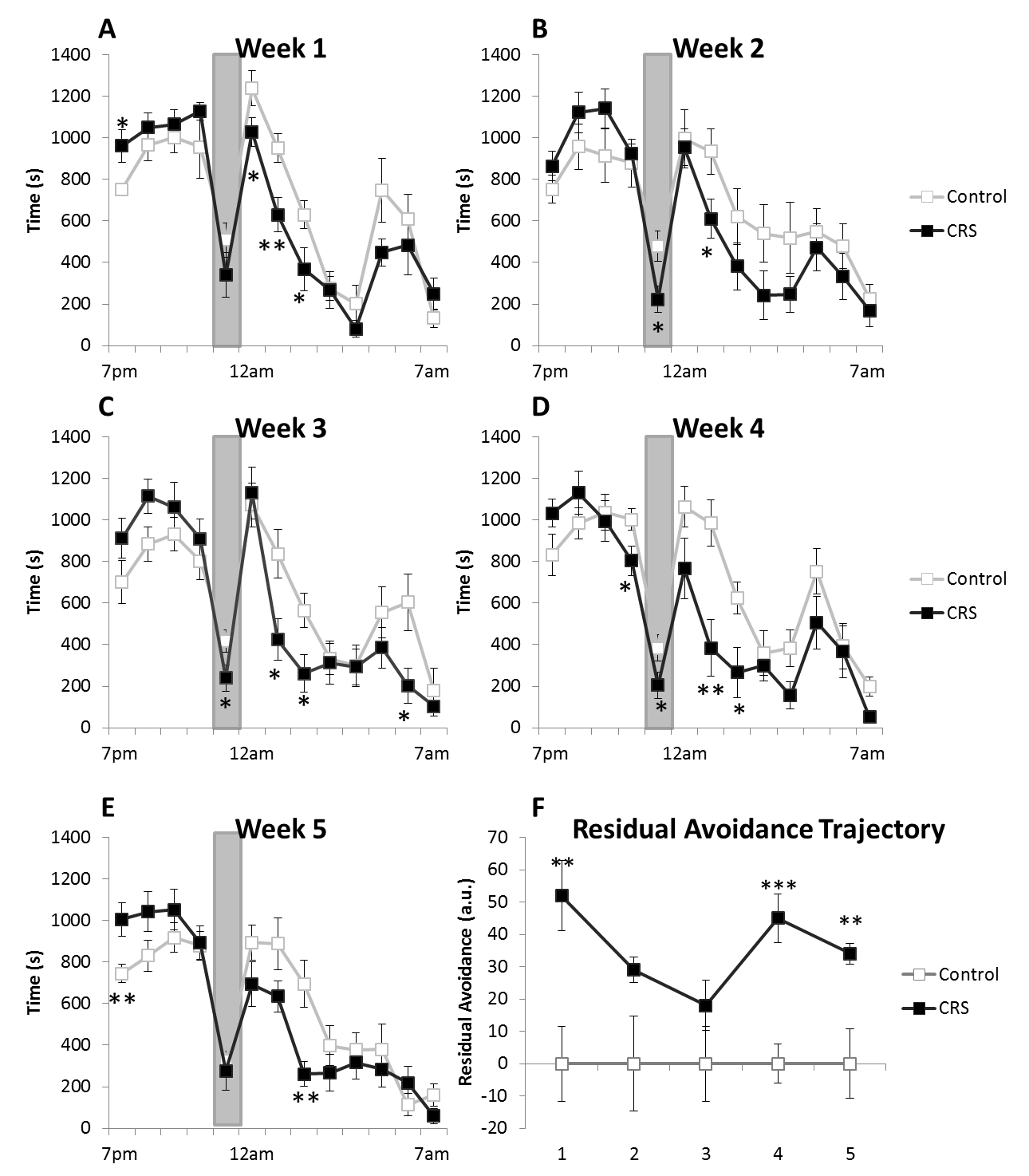

***Figure S5:* Chronic restraint stress induces behavioral alterations of the Food zone time in the PhenoTyper test**.

Mice were subjected to chronic restraint stress (CRS) for 5 weeks, and tested in the PhenoTyper test every week with a light challenge occurring between 11pm and 12am. The Food zone time was assessed every week (A-E). Using the residual avoidance (RA) calculation, each week’s food zone time was transformed into a single value resuming the trajectory of the effect of chronic stress exposure (F). *p < 0.05, **p < 0.01, ***p < 0.001 as compared with controls.

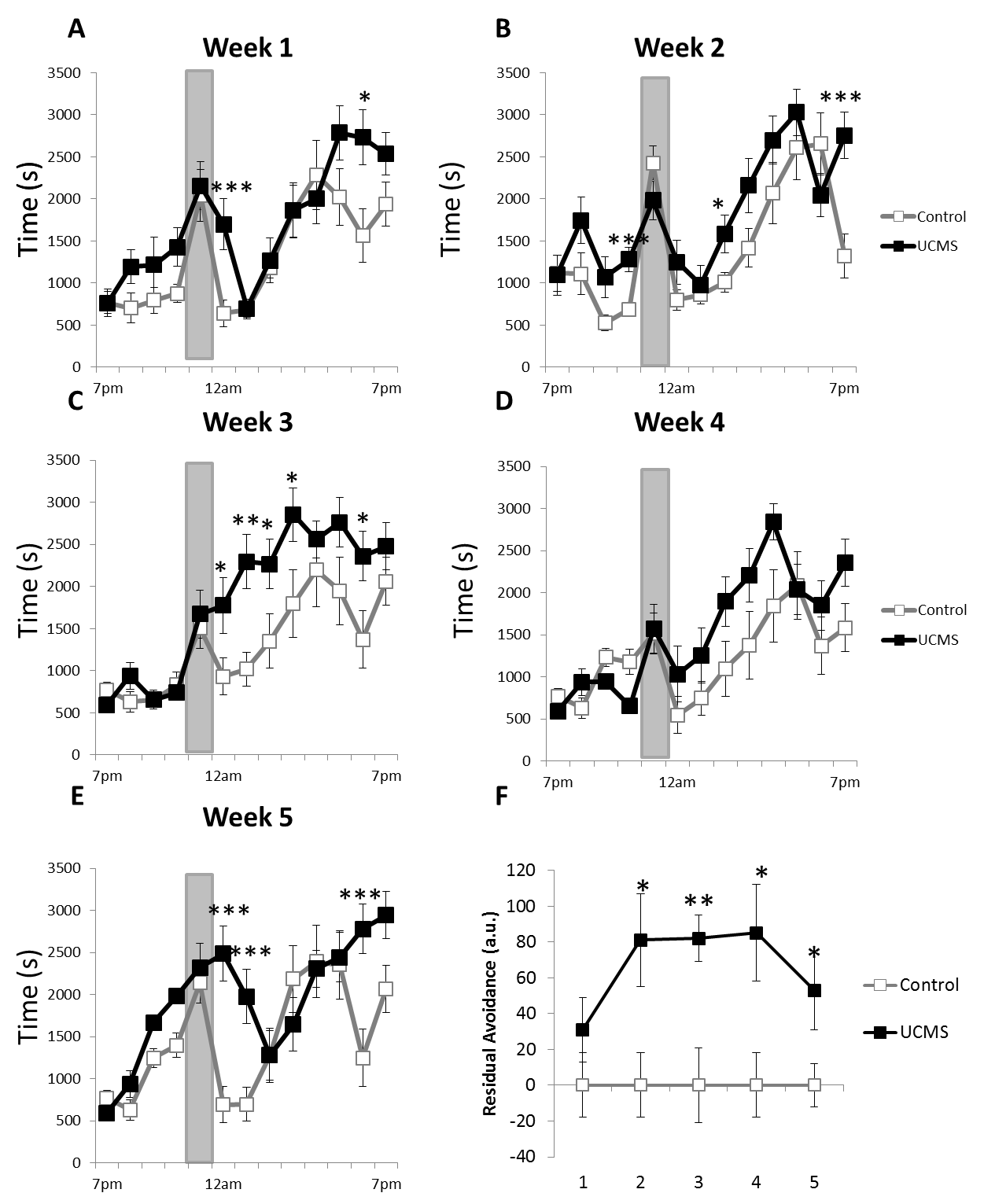

***Figure S6:* Unpredictable chronic mild stress induces behavioral alterations of the Shelter zone time in the PhenoTyper test**.

Mice were subjected to unpredictable chronic mild stress (UCMS) for 5 weeks, and tested in the PhenoTyper test every week with a light challenge occurring between 11pm and 12am. The Shelter zone time was assessed every week (A-E). Using the residual avoidance (RA) calculation, each week’s shelter zone time was transformed into a single value resuming the trajectory of the effect of chronic stress exposure (F). *p < 0.05, **p < 0.01, ***p < 0.001 as compared with controls.

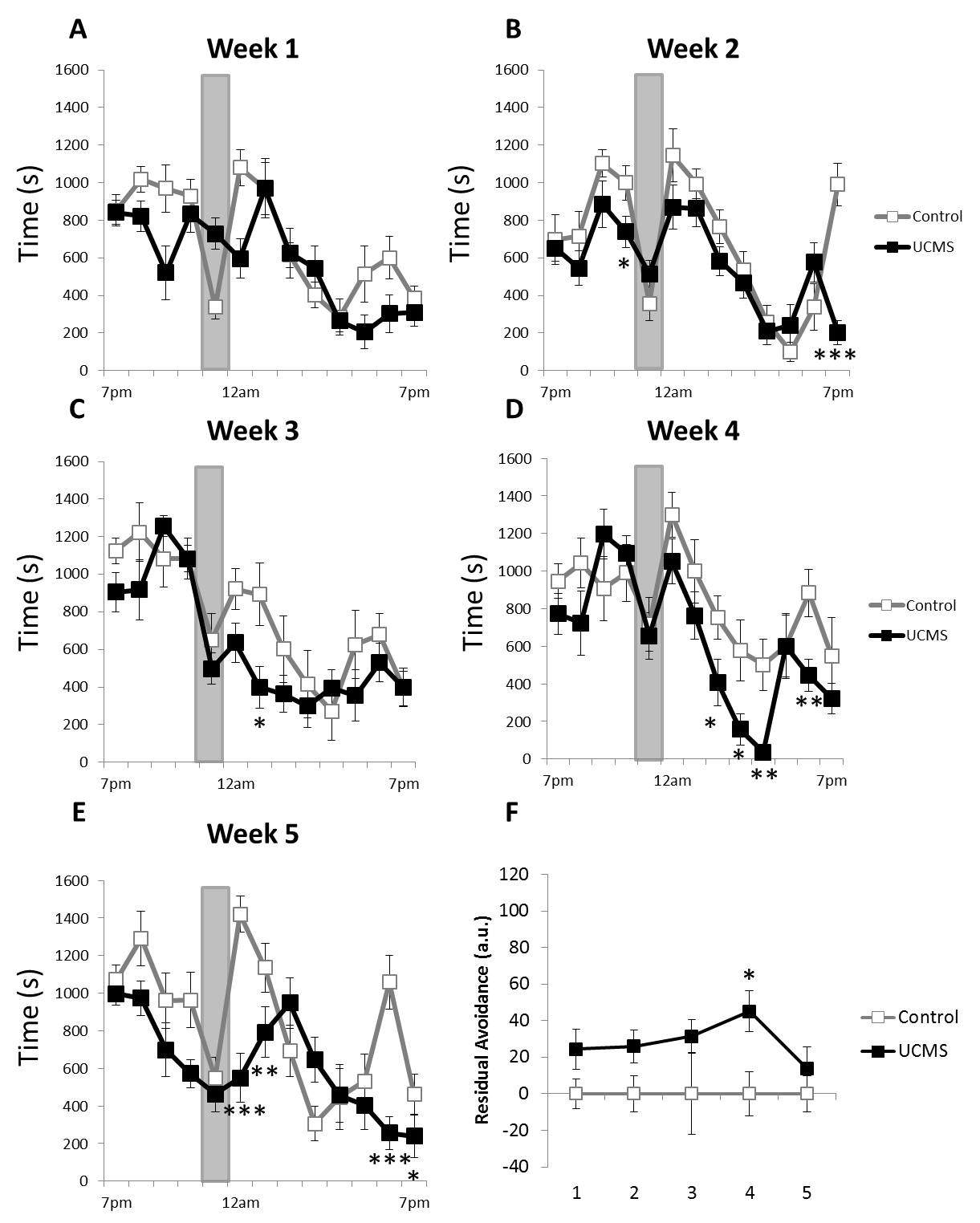

***Figure S7:* Unpredictable chronic mild stress induces behavioral alterations of the Food zone time in the PhenoTyper test**.

Mice were subjected to unpredictable chronic mild stress (UCMS) for 5 weeks, and tested in the PhenoTyper test every week with a light challenge occurring between 11pm and 12am. The Food zone time was assessed every week (A-E). Using the residual avoidance (RA) calculation, each week’s food zone time was transformed into a single value resuming the trajectory of the effect of chronic stress exposure (F). *p < 0.05, **p < 0.01, ***p < 0.001 as compared with controls.

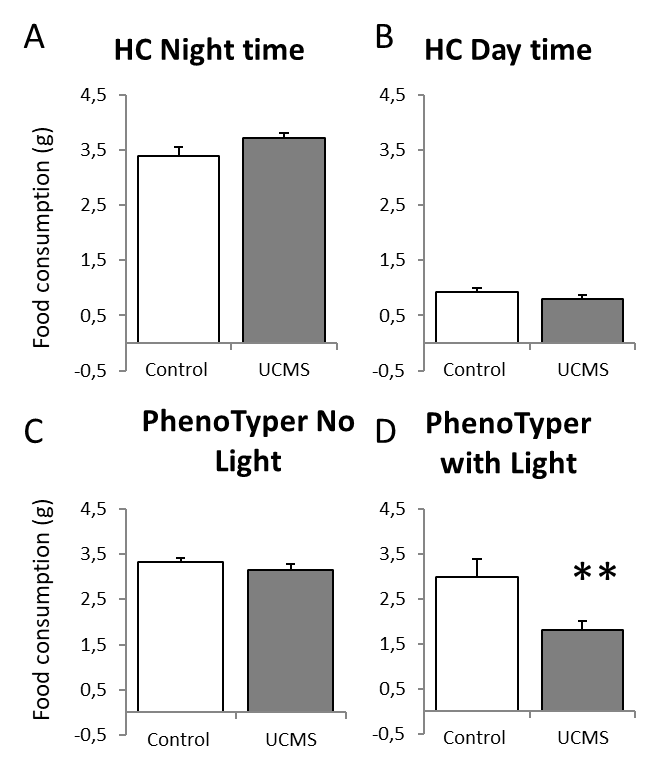

***Figure S8:* Food intake measurement in the PhenoTyper or in the animal home cage.**

In a separate cohort, food intake was measured in mice, in their home cage (A-B) or in the PhenoTyper boxes (C-D). Home cage food consumption during the dark cycle (A) or light cycle (B) was not statiscally different between control and UCMS mouse group (n=12/group, 50%females). In the PhenoTyper, if no light challenge was applied, UCMS and control mice showed similar total food intake (C). In the Phenotyper, application of the light challenge induced a significant difference between groups. Total food consumption was significantly reduced in UCMS-subjected mice (D). **p<0.01 compared to control.

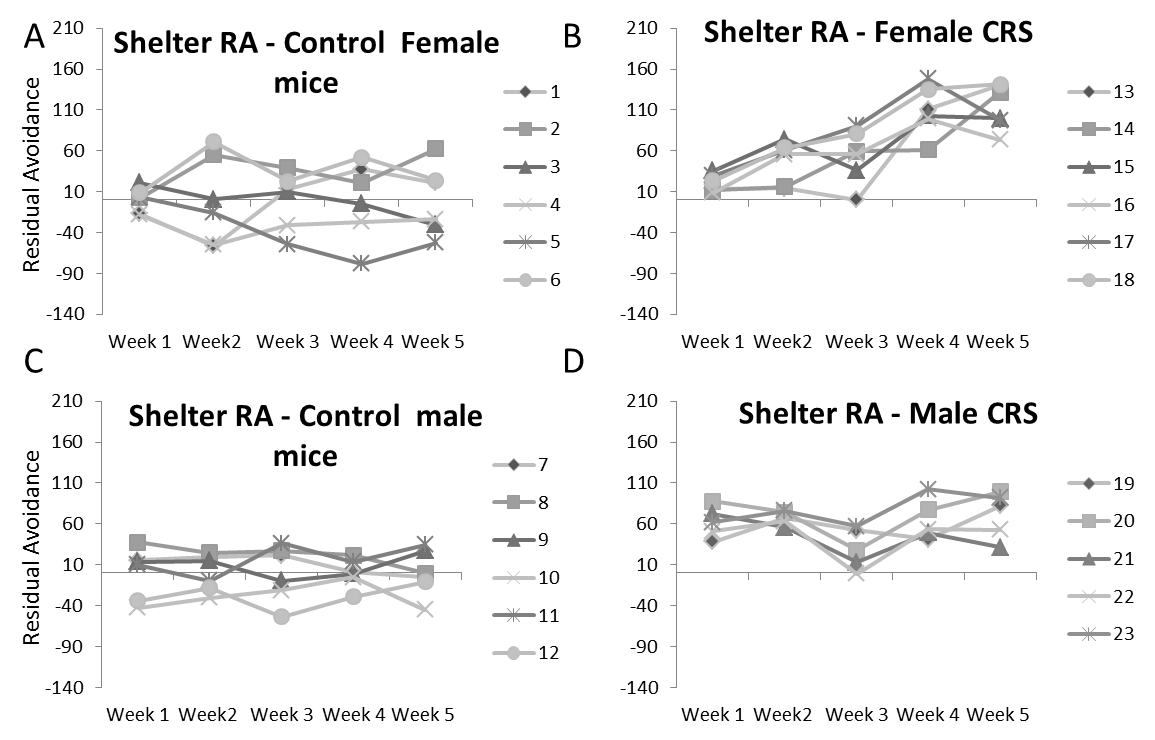

***Figure S9:* Individual trajectory of the residual avoidance measured in the shelter for each mouse, subjected to CRS or not.**

Residual avoidance (RA) was measured weekly in male and female mice control or subjected to CRS. Male and female control mice (A and C respectively) showed individual RA trajectory with limited variability between animals, and between weeks. Visualization of individual trajectory of CRS female mice (B) shows gradual increase in RA throughout the duration of the CRS protocol. Visualization of individual trajectory of UCMS male mice (D) shows increased RA throughout CRS exposure.

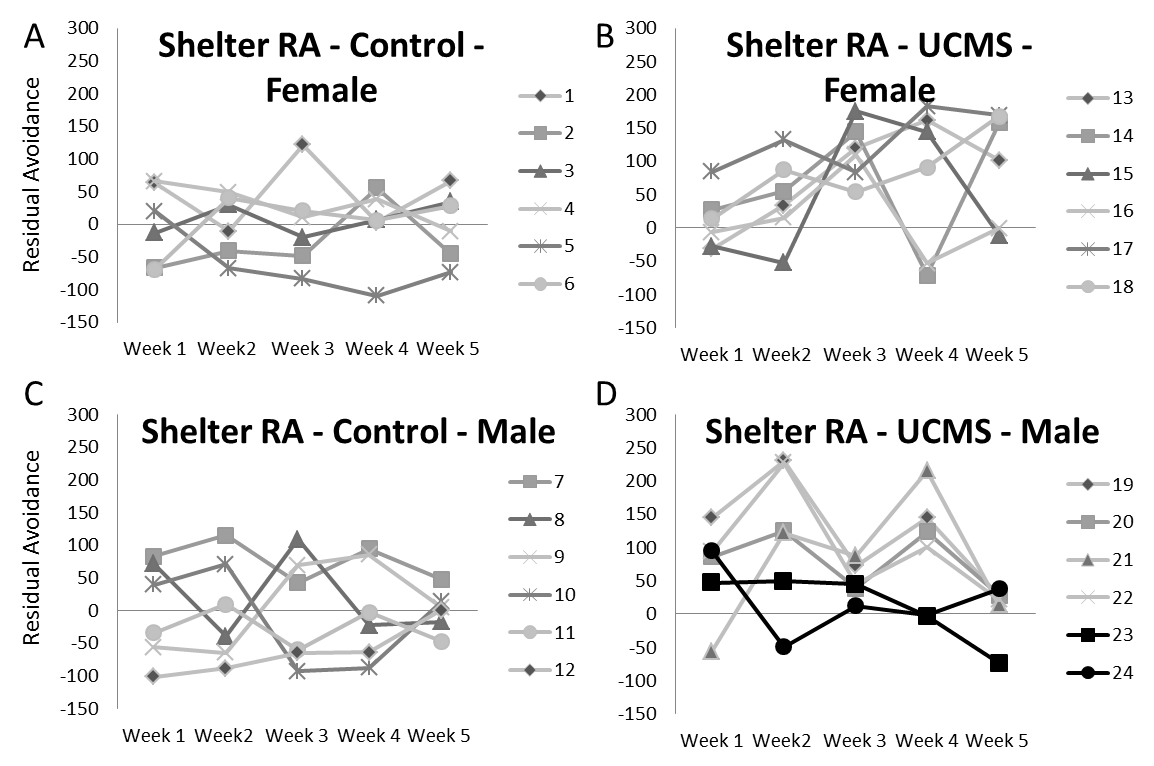

***Figure S10:* Individual trajectory of the residual avoidance measured in the shelter for each mouse, subjected to UCMS or not.**

Residual avoidance (RA) was measured weekly in male and female mice control or subjected to UCMS. Male and female control mice (A and C respectively) showed individual RA trajectory with variability between animals, but less within animals between weeks. Visualization of individual trajectory of UCMS female mice (B) shows that 3/6 showed gradually increasing RA throughout UCMS exposure and 3/6 showing RA increased for at least 2-3 time points throughout the UCMS exposure. Visualization of individual trajectory of UCMS male mice (D) shows that 2 of out 6 mice showed no change in RA (#23 and #24, in black) at any time points throughout the UCMS exposure.

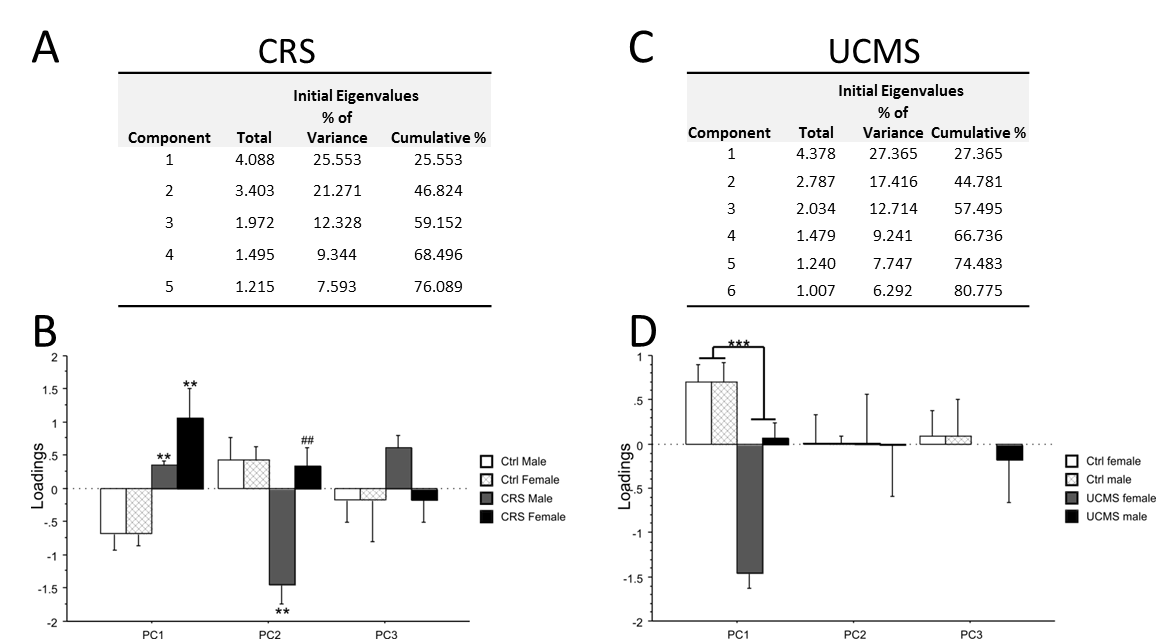

***Figure S11:*** **Principal component (PC) analysis of behavioral data obtained from mice exposed or not to chronic stress**

Initial eigenvalues (>1) representing the percentage variance in multiple principle components (PC) in chronic restraint stress (CRS, A) and unpredictable chronic mild stress (UCMS, C) cohorts. Analysis of the contribution of stress and sex on the 3 first components for CRS (B) and UCMS (D) experiments. **p < 0.01, ***p < 0.001 as compared with controls. ## p < 0.01 compared to males.
